# Triplex formation drives noncontiguous VIPR RNA-guided DNA recognition

**DOI:** 10.64898/2026.04.26.720927

**Authors:** Peter H. Yoon, Trevor A. Docter, Zeyuan Zhang, Kenneth Loi, Luis E. Valentin-Alvarado, Stephen G. Brohawn, Jennifer A. Doudna

## Abstract

Viral Interference Programmable Repeat (VIPR) systems use a noncontiguous code for RNA-guided transcriptional silencing. How the Vipr protein and a vrRNA comprising alternating GGY and NN segments achieve precise DNA targeting is unknown. Here we present 21 cryo-electron microscopy structures that span the VIPR assembly pathway. Vipr protomers oligomerize along the vrRNA to form a right-handed helical filament, sequestering each GGY motif and positioning the adjacent NN bases for target base pairing. DNA binding, in which every third nucleotide is skipped, results in target strand rotation to form a gapped vrRNA-DNA hybrid helix that wraps around the non-target DNA strand to form a structural triplex. These findings provide the structural basis of noncontiguous RNA-guided DNA binding in VIPR, establishing triplex-driven target-strand handoff as an elegant mechanism of programmable nucleic acid recognition.

## Introduction

RNA-guided systems solve nucleic acid recognition by coupling programmable base pairing to protein scaffolds. In every characterized instance, contiguous base pairing between the guide RNA and substrate dictates specificity (*1–4*). However, we recently discovered that bacteriophage-encoded Viral Interference Programmable Repeat (VIPR) systems use noncontiguous base pairing for RNA-guided DNA recognition (*5*). VIPR RNAs (vrRNAs) typically comprise 9-16 YNNGG tandem repeats that form alternating GGY and NN motifs. The NN dinucleotides encode specificity by pairing to the target DNA using a skip base rule in which a single nucleotide in the target is not paired between successive NN pairings. vrRNAs, in complex with the Vipr proteins homologous to CRISPR RAMPs (Cas5-7), direct transcriptional repression of competing phage genes (Fig. 1A). How this two component system employs this unexpected mode of nucleic acid recognition remains unknown.

**Figure 1.**
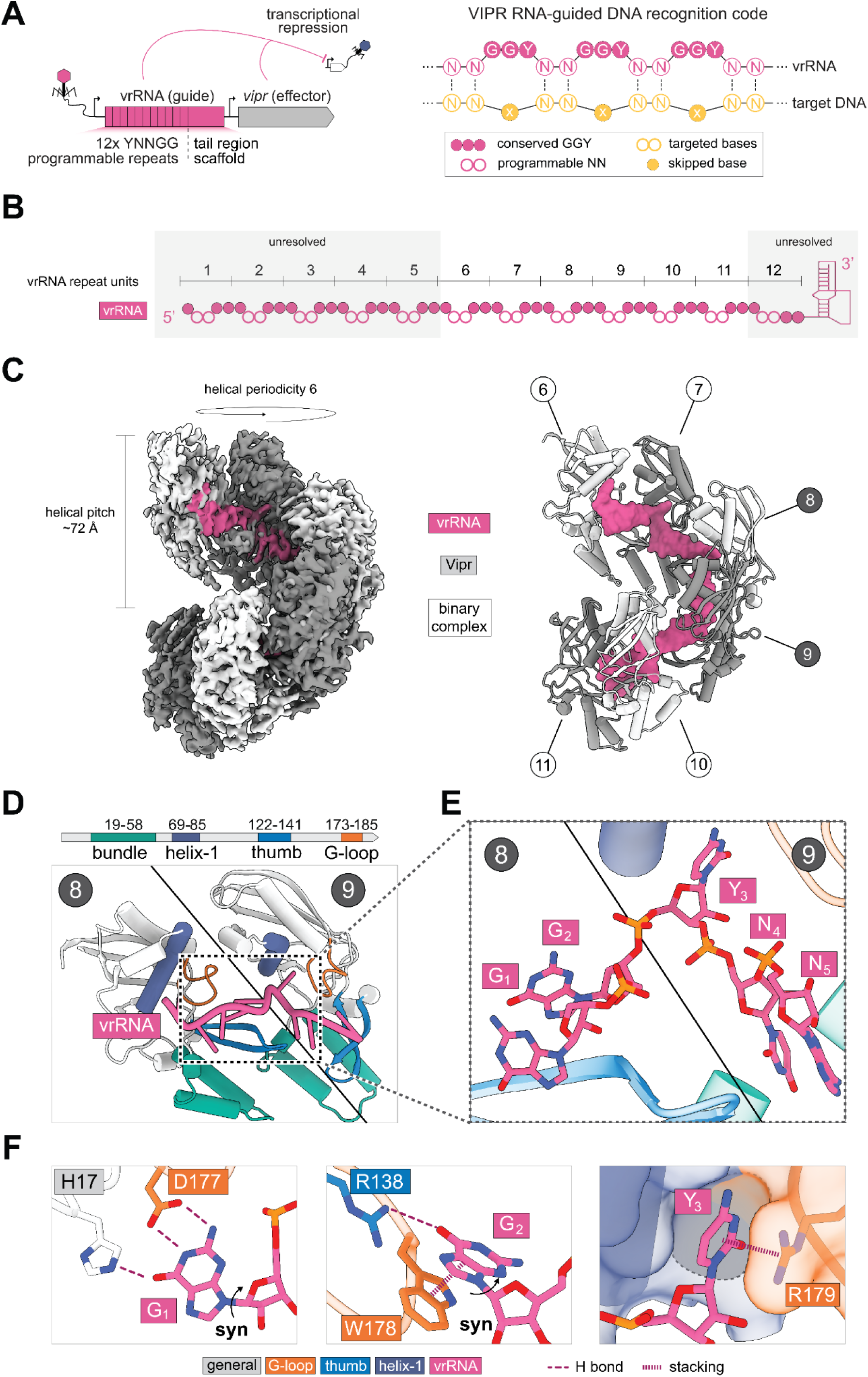
VIPR forms a helical assembly on vrRNA. **(A)** Cartoon representation of SUSP1 VIPR locus (left) and schematic of the GGY/NN motif-based RNA-guided DNA recognition (right). **(B)** Schematic of the 12-repeat vrRNA used for cryo-EM. Numbers label both the repeat unit and its bound Vipr protomer (e.g., subunit 10 = repeat 10). Color schematic same as in (A); regions not resolved in map are boxed in gray. **(C)** Cryo-EM map of the Vipr-vrRNA binary complex (bottom left), and cartoon model with alternating colors for Vipr protomer (bottom right). Circles with numbers inside correspond to the subunit number shown in the schematic. **(D)** Vipr protomer domain architecture (top) and interactions with vrRNA shown through two Vipr protomers spanning two vrRNA repeat units (bottom). **(E)** Zoom-in of GGY/NN motif interactions with Vipr. **(F)** Base-specific contacts at G_1_, G_2_, and Y_3_ positions. Hydrogen bonds (H bond) and stacking interactions indicated.

Here we present 21 cryo-electron microscopy (cryo-EM) structures of VIPR ribonucleoprotein (RNP) complexes that span their target recognition pathway from binary, to intermediate, and fully engaged ternary states. These structures reveal that Vipr proteins oligomerize along the vrRNA into a helical filament. Each Vipr protomer binds one repeat unit in the vrRNA, and adjacent protomers jointly displace the GGY bases to position the NN dinucleotides for gapped target pairing. Upon DNA binding, the VIPR filament constricts around the duplex. This strains the DNA into the helical register of the vrRNA to form an RNA-dsDNA triplex. Translocation of the complementary strand to enable base pairing with the vrRNA produces an R-loop triplex in which the non-target strand threaded through the R-loop core. This series of structures reveals the molecular basis of noncontiguous RNA-guided DNA recognition, implemented through triplex-driven strand exchange.

## Results

### Vipr proteins decode vrRNA repeats into a guide-presenting filament

Single-particle cryo-EM of Vipr-vrRNA complex (e.g., VIPR binary complex) purified from *E. coli* revealed that Vipr oligomerizes along the vrRNA as a right-handed helical filament in which one protomer is bound to one repeat (Fig. 1B; figs. S1-3; tables 1, 2). We resolved a monomeric class at 2.85 Å, and an additional dimeric class at 2.7 Å where two filaments interact at their 3’ ends (figs. S2, S3). Examination of the monomeric class reveals a helix with a six subunit periodicity and a ∼72 Å pitch that contains a central cleft housing the vrRNA (Fig. 1B, C). Although the sample contains 12 repeat units in the vrRNA (fig. S1), the map only showed modelable RNA and protein density for 6 repeats spanning the 6th to 11th repeat units, with weaker density at the 5th and 12th repeat units (Fig. 1B, C). This indicates the 5’ and 3’ ends are conformationally flexible in the binary complex.

Vipr proteins show the same RNA-binding mode seen in CRISPR-RAMPs. As in CRISPR-RAMPs (*6*), Vipr’s core RNA recognition motif (RRM) fold, glycine-rich loop (G-loop), and thumb domain all make extensive RNA contacts (Fig. 1D; fig S4). helix-1 of the RRM and the G-loop form the primary RNA-binding surfaces, while the thumb induces base-flipping at the GGY motifs critical for vrRNA recognition by Vipr. These shared architectural and mechanistic features are consistent with the proposed evolutionary relationship between VIPR and CRISPR RAMPs (*5*).

Two adjacent Vipr protomers jointly displace one GGY (G_1_, G_2_, and Y_3_) motif to position the NN (N_4_, N_5_) dinucleotides for gapped target pairing (Fig. 1D, E). Within the GGY bases, the guanines adopt the syn glycosidic conformation and present their Watson-Crick face to the protein. Within the first protomer, G_1_ is recognized by hydrogen bonds from the RRM β-sheet (H17) and the G-loop (D177), whereas G_2_ is cradled between the thumb (salt bridge, R138) and G-loop (π-stacking, W178) (Fig. 1F). Alanine substitutions in D177 or R138 abolished VIPR-mediated green fluorescent protein (GFP) silencing in *E. coli*, consistent with the functional importance of these residues (fig. S5). Within the second protomer, Y_3_ is held in a narrow pocket formed by helix-1 (steric restriction, E71, M72) and the G-loop (cation-π, R179, steric restriction, C173, T175) that sterically excludes purines. These contacts lock the GGY bases, allowing specificity to be dictated solely by the programmable NN dinucleotides.

### A gapped code for RNA-guided DNA recognition

Single-particle cryo-EM of VIPR binary complex incubated with a pre-unwound dsDNA target (e.g., paired on flanks, unpaired in center) revealed a constricted helical filament that threads both DNA strands through its core (Fig. 2A, B; figs. S6, S7; tables 1, 2). We observed a complete class at 3.13 Å that resolved all twelve Vipr subunits and the terminal pseudoknot that caps the 3’ end of the RNA (Fig. 2A, B; fig. S8). We also observed an alternative class at 3.17 Å that did not resolve the 5’ most repeat unit, but was otherwise identical (fig. S7). The ternary filaments are constricted along their entire length with a reduced periodicity (five vs. six subunits) and helical pitch (∼66 Å vs. ∼72 Å), sealing the gap between successive turns found in the binary state (Figs. 1C and 2B, C, Movie 1).

**Figure 2.**
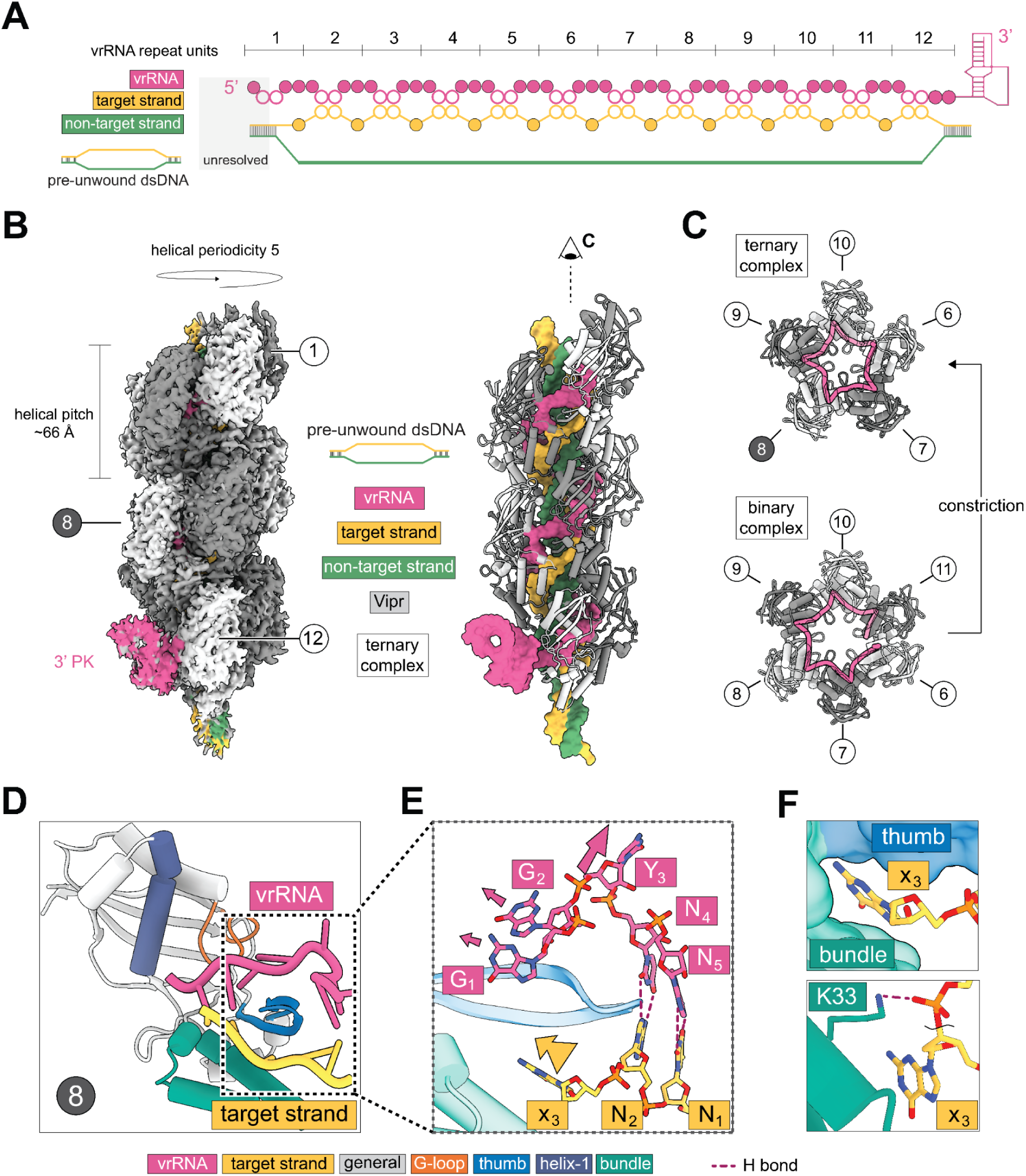
VIPR code for gapped RNA-programmed DNA recognition. **(A)** Schematic of the Vipr-vrRNA-DNA ternary complex with pre-unwound dsDNA substrate and its register against the vrRNA across all 12 Vipr protomers. **(B)** Cryo-EM map of the Vipr-vrRNA-DNA ternary complex with 3’ pseudoknot (PK) (left). Cartoon model colored by component (right). **(C)** Top-down views of ternary (top) and binary (bottom) complexes. **(D)** Vipr protomer interactions with vrRNA and target strand DNA. **(E)** Zoom-in of gapped R-loop base pairing between vrRNA (pink) and target strand (yellow). **(F)** Accommodation of the target strand x_3_ skipped base by the thumb (top) and bundle (bottom) domains. H bonds indicated.

The central cleft of the constricted filament contains the vrRNA-DNA gapped duplex. (Fig. 2D, E). This unusual R-loop structure is achieved by Vipr protein thumb-domain insertions that periodically break the guide-target duplex to asymmetrically displace nucleotides on both strands (Fig. 2A, E). On the guide, the thumb displaces all three GGY bases (G_1_, G_2_, and Y_3_), while on the target only the skipped base (x_3_) is displaced. This enables pairing between the NN on the guide (N_4_ and N_5_) to the matching NN on the target (N_1_ and N_2_), defining the VIPR non-contiguous recognition code. Notably, the Y_3_ on the guide and the x_3_ skip base on the target point in opposite directions, which is reminiscent of the antiparallel base-flip geometry observed in Class 1 CRISPR effector complexes (Fig. 2D, E; fig. S9) (*7*).

The extruded x_3_ skip base is accommodated by a narrow pocket formed between the α-helical bundle and the thumb that accommodates one nucleotide (Fig. 2F). The α-bundle additionally contacts the phosphate backbone adjacent to the skip base (K33), and mutation of this residue to alanine abolished VIPR-mediated GFP repression in *E. coli* (fig. S5). An analogous interaction is found in Class 1 CRISPR, where the small subunit similarly composed of a helical bundle stabilizes a single flipped target base against the RAMP thumb domain (fig. S9) (*7*). The gapped VIPR code thus arises from a thumb-bundle module shared with Class 1 CRISPR, despite encoding fundamentally different pairing rules.

### VIPR target engagement results in an R-loop triplex

In addition to the gapped RNA-DNA duplex, the ternary complex core surprisingly houses the non-target strand that threads through the duplex to form a geometric triplex in which all three strands share a common helical pitch (Fig. 3A). We call this unique structure the R-loop triplex (Fig. 3A). Although all three strands have a helical pitch of ∼66 Å, this is achieved with different numbers of nucleotides per turn between the RNA and the DNA. The vrRNA contributes ∼25 nucleotides per turn, while both the target and non-target DNA strands contribute 15 (Fig. 3B). This owes the asymmetrical geometry of the gapped R-loop (e.g., 5 bases on RNA per 3 on target DNA), which forces the vrRNA backbone to traverse a longer path than either DNA strand to maintain the shared helical pitch.

**Figure 3.**
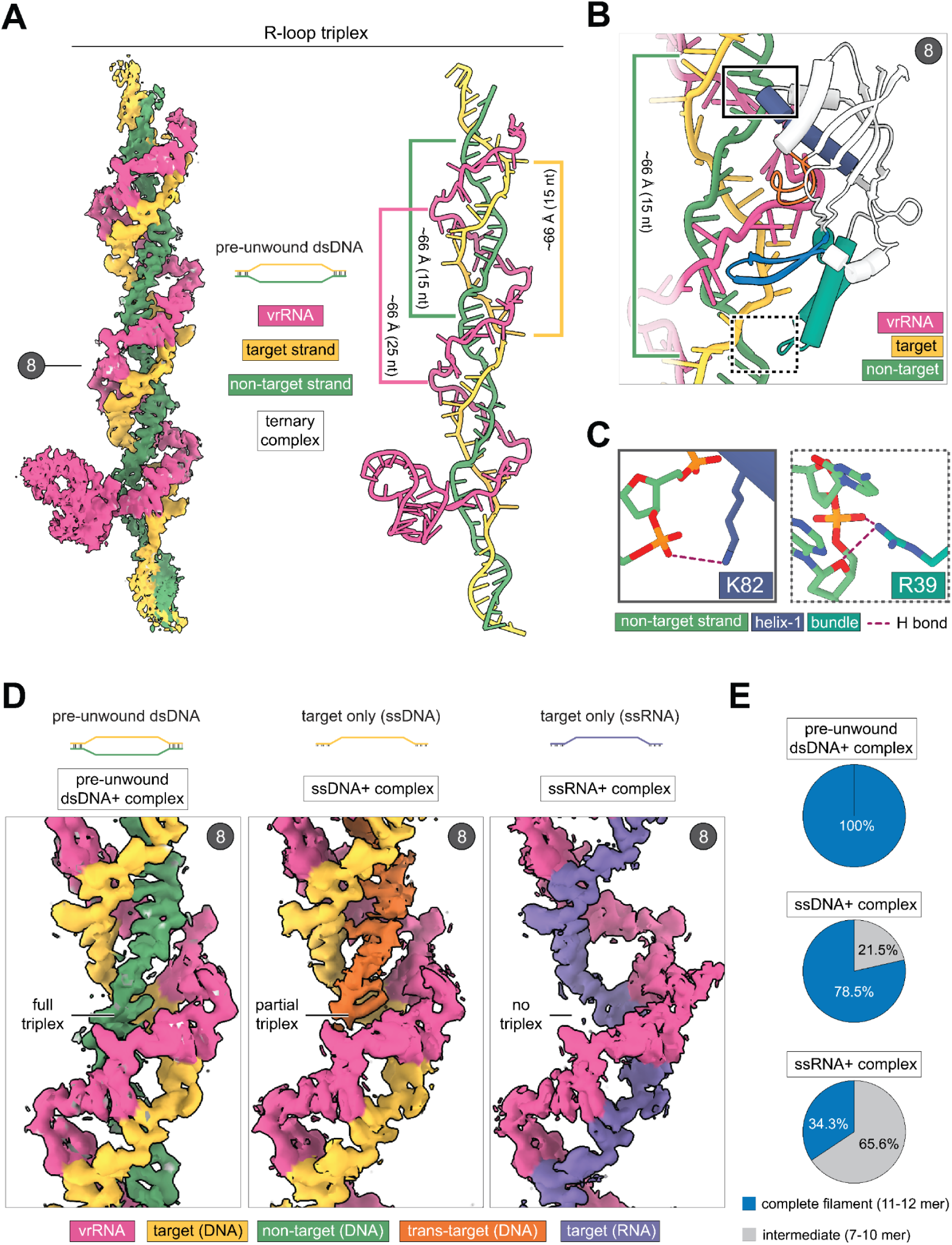
VIPR targeting state forms a geometric triplex of nucleic acids. **(A)** Carved cryo-EM density of the R-loop triplex in the pre-unwound dsDNA ternary complex (left). Model with segment lengths indicated (right). **(B)** Vipr protomer interactions at the R-loop junction with vrRNA, target strand, and non-target strand. **(C)** Non-target strand contacts by helix-1 (K82, left) and bundle (R39, right) domains. **(D)** Carved cryo-EM densities of complexes assembled with dsDNA, ssDNA, or ssRNA targets. **(E)** Distribution of complete filament (11-12 subunits) versus intermediate (7-10 subunits) particles for each substrate.

Unlike previously reported triplexes held together by inter-strand contacts (*8*, *9*), the R-loop triplex is held in place by the Vipr proteins as the non-target strand makes no contacts with the R-loop. Each Vipr protomer clamps the non-target strand at two points, one at each end of the subunit (Fig. 3B). The first clamp is formed by the top tip of helix-1, which engages the non-target strand backbone through hydrogen bonds (K82, K88) and amide interactions (G87, K88) (Fig. 3C, left; fig. S10). The second clamp is formed by the bottom tip of the α-bundle (R39) that makes similar interactions (Fig. 3C, right). Together, these top-and-bottom clamps fix each subunit’s spacing along the non-target strand, enforcing the ∼66 Å pitch shared across all three strands. Alanine substitution of K82 abolishes and R39 reduces VIPR-mediated repression in *E. coli* (fig. S5), supporting an essential role for this non-target strand engagement surface.

Cryo-EM of VIPR binary complex incubated with ssDNA or ssRNA revealed that the filament will recruit a second strand in trans to reconstitute the triplex (figs. S11-S14; tables 1, 3). Incubation of VIPR binary complex with only the target strand (ssDNA) resulted in three distinct classes that resolved 10, 11, and 12 subunits (3.08 Å, 3.00 Å and 3.03 Å; figs. S11, S12; table 1, 3). In every class, the ssDNA complex formed the R-loop triplex by incorporating a second copy of the target strand in trans (Fig. 3D). In contrast, incubation with an RNA analog of the target strand (ssRNA) yielded two classes with 11 and 7 subunits (2.76 Å and 2.96 Å; figs. S13, S14). Both classes formed a vrRNA-RNA duplex, but showed no density for a third strand (Fig. 3D). VIPR thus accommodates both DNA and RNA in the guide-target duplex, but triplex formation requires the third strand to be DNA.

Particle distributions across the three substrates suggest the R-loop triplex acts to stabilize the filament (Fig. 3E, S15). The pre-unwound dsDNA substrate resulted exclusively in complete complexes (11-12 subunits) in which all clearly resolved NN dinucleotides engaged the target. The ssDNA substrate produced mostly complete complexes (79%) with some stalling as incomplete 10-subunit intermediates (20%). In contrast, the ssRNA substrate that cannot form the triplex shifted dramatically toward incomplete assembly. The majority of particles produced a 7-subunit intermediate (66%), and only a minority reached the 11-subunit state (34%). The relevance of the triplex is further supported by structures of *Pseudomonas fulva* prophage (*P. fulva*) VIPR ternary complexes, which show a conserved R-loop triplex architecture (fig. S16-19, tables 1,4). The R-loop triplex is therefore a general feature of VIPR systems that appears to be essential for both filament completion and targeted transcriptional silencing.

### Target strand handoff converts RNA-dsDNA triplex into R-loop triplex

Cryo-EM of VIPR binary complex incubated with normal dsDNA (e.g., without pre-unwinding) revealed a continuum of assembly intermediates that provide snapshots of DNA unwinding (Fig. 4A; figs. S20-24; tables 1, 5). These intermediates ranged from 5 to 11 Vipr subunits in complex with progressively unwound DNA. All classes showed clear density for subunits 8-12 near the 3’ pseudoknot, and additional protomers resolved in an apparent 3’-to-5’ direction along the vrRNA (Fig. 4A, top, Movie 2). Across these intermediates, the appearance of additional protomers runs ahead of R-loop formation. In three transitions (6-to-7, 7-to-8, and 9-to-10), additional 5’ protomers and their vrRNA segments resolve without corresponding gains in vrRNA-target pairing (Fig. 4A).

**Figure 4.**
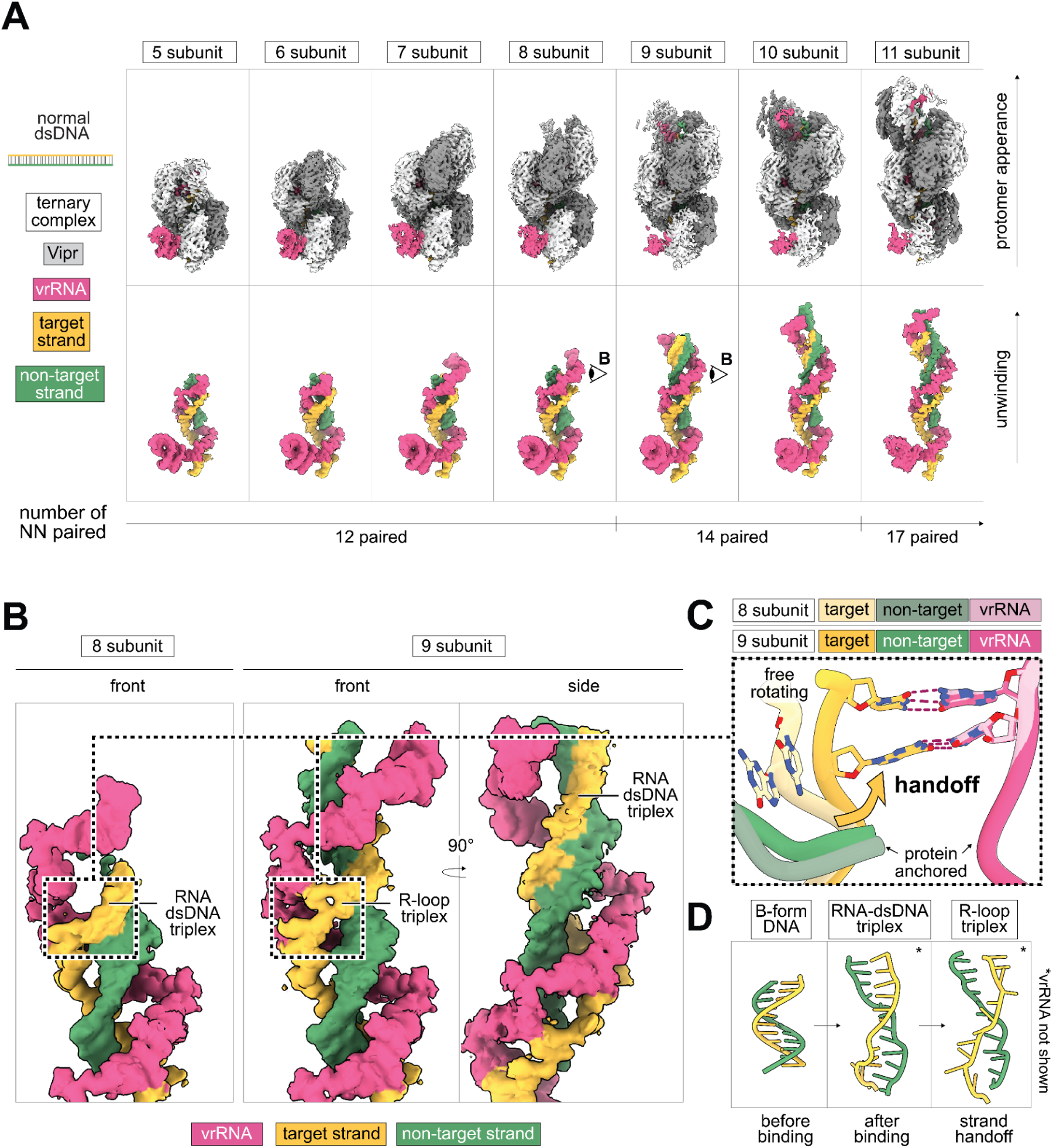
VIPR unwinds DNA via triplex mediated handoff mechanism. **(A)** Density maps of the VIPR complex from 5 subunit to 11 subunit ternary complex states (top) and the carved density for associated nucleic acid assemblies (bottom), arranged along the unwinding reaction coordinate. **(B)** Close up of the 8-subunit (front) and 9-subunit (front and side) states comparing leading edge RNA-dsDNA triplex and R-loop triplex. **(C)** Focused view of target base hand off from the non-target strand (8 subunits) to vrRNA (9 subunits). **(D)** DNA models depicting progression from B-form to RNA-dsDNA triplex (normal dsDNA, 10-subunit) to R-loop triplex (pre-unwound dsDNA, 12-subunit) driven by Vipr binding.

Further investigation of the 8-to-9 subunit transition reveals how progressive DNA unwinding is coupled to protomer addition (Fig. 4B). At the leading 5’ edge of both the 8 subunit and 9 subunit states, the newly resolved protomer forms an RNA-dsDNA triplex where a strained dsDNA is held in the vrRNA’s helical register as seen in the R-loop triplex state (Fig. 4B, boxed regions). In this intermediate state, the non-target strand is already stabilized by the same protomer interface used in the R-loop triplex (fig. S10). In transitioning from 8-to-9 subunits, the RNA-dsDNA triplex region of the 8 subunit state resolves into an R-loop triplex in the 9 subunit state where the target strand is now paired to the vrRNA (Fig. 4B). This supports the RNA-dsDNA triplex as a structural precursor to R-loop formation. Consistent with this, a side view of the 9-subunit class shows that the leading edge has advanced, with a new RNA-dsDNA triplex now positioned ahead of the nascent R-loop triplex region (Movie 3).

Overlay of the 8– and 9-subunit states reveals that conversion to an R-loop triplex proceeds through a local target strand handoff (Fig. 4C, Movie 4). Between the two states, the target strand bases rotate from the non-target strand face to instead pair with the vrRNA. The latter two remain largely stationary, consistent with them being anchored by the Vipr protein within the constricted VIPR filament. This handoff can occur independently at each position, as seen in the 11-subunit complex where only the leading edge has completed the handoff and the middle segment remains an RNA-dsDNA triplex (Fig. 4A; fig. S25; Movie 4). This unconverted middle region is identical to the equivalent region in the 10-subunit state (Fig. 4A; figs. S24), implying localized target strand movement converts an RNA-dsDNA triplex into an R-loop triplex. Together, these observations support a model of triplex-driven DNA unwinding in which Vipr protein engagement with target DNA builds a triplex scaffold, and local target strand handoff completes the R-loop at each position (Fig. 4D).

## Discussion

The series of 21 cryo-EM structures of the SUSP1 and *P. fulva* VIPR complexes presented here reveal the molecular basis of VIPR guide RNA coordination, target engagement, and sequence interrogation. Vipr protomers oligomerize along the vrRNA by binding to the GGY motifs, forming a relaxed filament that positions the NN dinucleotides for base pairing with the target. This filament binds to and constricts the target DNA duplex, mediating sequential and directional unwinding. Along this trajectory, the target duplex persists alongside the vrRNA to form an RNA-dsDNA triplex. This is superseded by a final R-loop triplex where the target strand separates from the non-target strand to instead pair with the vrRNA via gapped recognition.

Two protein domains, the thumb and the α-bundle enable gapped base pairing by VIPR. The thumb displaces the GGY bases of the vrRNA and the skipped base on the target strand to enable basepairing only by the flanking dinucleotides. In Class 1 CRISPR, RAMPs similarly contain a thumb domain that periodically displaces every sixth base-pair between the guide and target strands. Cas11 small subunits cradle the flipped target-strand base similarly to the α-bundle. In Class 2 CRISPR systems, a single domain in the ancestral effector that mediates multiple functions is proposed to have separated into multiple domains with specialized functions (*10*). These parallels raise the possibility that Class 1 effector complexes emerged through specialization of modules already present in a Vipr-like progenitor.

Despite architectural parallels to CRISPR, the VIPR unwinding mechanism is distinct from known RNA-guided DNA-targeting systems that unwind DNA through base-pairing-driven R-loop formation (*11–15*). VIPR structures instead suggest that protein-driven nucleic acid triplex formation rather than RNA invasion of the duplex, facilitates DNA unwinding to enable RNA-DNA base pairing. Protomers at the leading edge contact the non-target strand of the still-unseparated duplex, capturing it into an RNA-dsDNA triplex that distorts and destabilizes pairing between the DNA strands. With the non-target strand and vrRNA anchored by the protein, the target strand alone dissociates and pairs with the vrRNA to yield the final R-loop triplex. This distinct mechanism may be required because a gapped guide provides only partial complementarity at any register and is presumably insufficient to thermodynamically outcompete the DNA-DNA duplex.

VIPR represents a programmable platform for nucleic acid targeting with unprecedented flexibility. The Vipr protein makes no base-specific contacts with the target DNA, and sequence specificity is entirely mediated by the vrRNA guide. The structures further imply that vrRNA length dictates VIPR filament length such that both the target sequence and footprint are programmable. Consistent with this, natural VIPR systems show variable vrRNA lengths, with some reaching up to 30 repeats, implying that long stretches of DNA can be covered by a single VIPR complex. VIPR systems are further diversified through their associated accessory genes, which include predicted helicases and nucleases. Whether these accessory genes sense the unusual nucleic acid geometry or protein filament conformational changes caused by VIPR target recognition remains unclear, but their characterization could unlock another layer to sequence programmable biotechnologies.

## Acknowledgements

We thank Jamie Cate for extensive discussions and help with manuscript editing; members of the Doudna, Cate, and Brohawn labs for technical advice and insightful discussions. HHMI covered open access publication charges.

## Author contributions

**Conceptualization: PHY, TAD, ZZ, JAD**

**Investigation: PHY, TAD, ZZ, KJL, LAV**

**Visualization: PHY, TAD, ZZ**

**Funding acquisition: JAD**

**Supervision: JAD, SGB**

**Writing – original draft: PHY, TAD, ZZ, JAD**

**Writing – review & editing: PHY, TAD, ZZ, KJL, LAV, SGB, JAD**

## Data, code, and materials availability

All data, code, and materials used in the analysis will be made publicly available online.

## Supplementary Materials

Figs. S1 to S25

Tables S1 to S5

Movies S1 to S4

## Supplementary Materials for

**The PDF file includes:**

Materials and Methods

Figs. S1 to S25

Tables S1 to S5

## Materials and Methods

### Cloning, expression and purification

SUSP1 and *P. fulva* VIPR RNPs were expressed and purified as described in the VIPR discovery paper (*10*). Briefly, VIPR was expressed in *E. coli* from pJEX-SC101 (SUSP1, vrRNA1 and vipr, C-terminal 6xHis on Vipr) or pBAD-ColE1 (*P. fulva*, vrRNA1–3, vipr, and vap1, C-terminal Twin-Strep on Vipr). SUSP1 RNPs were captured on a Ni-NTA Superflow cartridge (Qiagen) in 50 mM Tris pH 8.5, 500 mM NaCl, 5 mM MgCl2, 1 mM TCEP, whereas *P. fulva* RNPs were captured on Strep-Tactin-XT (Cytiva) in 50 mM Tris pH 8.5, 300 mM NaCl, 5 mM MgCl₂, 1 mM TCEP and eluted by on-column TEV protease cleavage. Both RNPs were then purified by Heparin chromatography (Cytiva) and resolved on a Superose 6 Increase column (Cytiva) in 50 mM HEPES pH 6.8, 100 mM KCl, 5 mM MgCl2, 0.1% glycerol, 1 mM TCEP, which was also used as the cryo-EM buffer. The *P. fulva* RNP co-purified with a bound DNA substrate, presumably an off-target site in 10-beta *E. coli* Mix & Go! Competent Cells (Zymo) used in expression, and this endogenously captured ternary complex was used directly for cryo-EM without addition of exogenous DNA.

### Cryo-EM Sample Preparation and Grid Freezing for Cryo-EM

For all SUSP1 constructs, a 3.0 μL sample at the concentrations noted in table 1 was applied to UltrAuFoil 300 mesh R 1.2/1.3 gold grids (Quantifoil, Großlöbichau, Germany) that were freshly glow discharged at 25mA for 25s on 10s prep. Sample was incubated for 5 seconds at 4°C and 100% humidity prior to blotting with Whatman #1 filter paper for 3 seconds at blot force 1 and plunge-freezing in liquid ethane cooled by liquid nitrogen using a FEI Mark IV Vitrobot (FEI/Thermo Scientific, USA). All target substrates were added to binary VIPR at a 2-fold molar excess and left to incubate at room temperature for 30 minutes prior to blotting and vitrification.

For the SUSP1 binary complex, grids were clipped and stored in liquid nitrogen. Data was collected on a Titan Krios G3i electron microscope operated at 300 kV. Dose-fractionated images (50 electrons per Å2 over 50 frames) were recorded on a K3 direct electron detector (Gatan) in regular resolution counting mode with pixel size of 0.848 Å. 363 movies were collected in a 11 × 11-hole pattern with three targets per hole around a central hole position using image shift. Defocus was varied from −0.6 to −1.6 μm using SerialEM (*16*). See Table 2 for data collection statistics.

For all SUSP1 ternary datasets, grids were clipped and stored in liquid nitrogen. Data was collected on an FEI Talos Arctica electron microscope operated at 200kV. Dose-fractionated images (50 electrons per Å^2^ over 50 frames) were recorded on a K3 direct electron detector (Gatan, USA) in regular resolution counting mode with a pixel size of 1.14 Å. Movies were collected in a 7 × 7-hole pattern with one shot per hole around a central hole position using image shift. Defocus was varied from −0.8 to −1.8 μm using SerialEM. See Tables 2-4 for data collection statistics.

For our *P. fulva* data sample, A 3.5 μL sample at 4 µM was applied to UltrAuFoil 300 mesh R 1.2/1.3 gold grids coating (Quantifoil, Großlöbichau, Germany) that were freshly glow discharged at 25mA for 25s on 10s hold. Sample was incubated for 5 seconds at 4°C and 100% humidity prior to blotting with Whatman #1 filter paper for 3 seconds at blot force 1 and plunge-freezing in liquid ethane cooled by liquid nitrogen using a FEI Mark IV Vitrobot (FEI/Thermo Scientific, USA).

For the *P. fulva* complex, grids were clipped and stored in liquid nitrogen. Two datasets were separately collected on a Titan Krios G2i electron microscope operated at 300 kV. Dose-fractionated images (50 electrons per Å2 over 50 frames) were recorded on a K3 direct electron detector (Gatan) in regular resolution counting mode with pixel size of 0.9432 Å. 121 movies were collected in a 11 × 11-hole pattern with one target per hole around a central hole position using image shift. Defocus was varied from −0.6 to −1.6 μm using SerialEM. See Table 5 for data collection statistics.

### Cryo-EM Data Processing and Data Collection

For the SUSP1 VIPR binary complex, 6174 micrographs were motion-corrected with patch motion-correction and contrast transfer function parameters were fit with patch CTF in cryoSPARC v4.6.2 (*17*). 5,472,885 particles were selected by blob picking on micrographs CTF fit to 5.0 Å or better. One round of 2D classification yielded 2,017,019 particles which were used for a 3-class ab initio. One class with protein-like density was selected for 971,459 particles. These particles were subjected to one 2-class ab initio (maximum resolution of 3 Å, initial resolution of 10 Å, a Fourier step radius of 0.01, center structures in real space set to false, and a class similarity of 0. We note that these settings were used for all other ab initio jobs unless clearly stated otherwise), one round of 3D classification in which we selected for classes with the highest resolution, and one additional 2-class ab initio. Non-uniform refinement of the higher quality map yielded a consensus map at an overall resolution of 2.68 Å with 462,803 particles. Interestingly, this map showed an amorphous density toward the 3’ end of the complex that was only visible at low-contour. We investigated this by re-extracting our particles at both box sizes of approximately 510 Å and 680 Å. Both sets were used for 2-class ab initios to determine which would give the higher resolution.

In both cases, we resolved two separate classes: a monomeric class and a dimeric class, where two binary complexes form a weak interaction at a 3’ pseudoknot density. Non-uniform refinement of the 510 Å box-generated monomeric class yielded a map at an overall resolution of 2.85 Å with 213,691 particles. For the dimeric class, density at the 3’ pseudoknot was quite weak in the 510 Å box-generated class compared to the 680 Å box-generated class, while the overall resolution within the 510 Å box-generated class was demonstrably higher. As such, we used the map from the 680 Å box-generated map to make a mask in ChimeraX as it had the strongest density for the pseudoknot. We used this mask to recenter the 510 Å box dimeric particles around the pseudoknot interaction between the two complexes. A final ab initio and non-uniform refinement yielded a map with an overall resolution of 2.70 Å with 212,926 particles with improved resolution at the 3’ pseudoknot without a dramatic loss in resolution at the more distal 5’ subunits.

For SUSP1 VIPR incubated with a pre-unwound dsDNA target, 5,253 micrographs were motion-corrected with patch motion-correction and contrast transfer function parameters were fit with patch CTF in cryoSPARC v4.6.2. 3,903,238 particles were selected by blob picking on micrographs CTF fit to 6.0 Å or better. One round of 2D classification yielded 1,129,858 particles which were used for a 3-class ab initio. Two classes with protein-like density were selected for 875,987 particles which were then used for an additional 3-class ab initio (maximum resolution of 3 Å, initial resolution of 10 Å, a Fourier step radius of 0.01, center structures in real space set to false, and a class similarity of 0. We note that these settings were used for all other ab initio jobs unless clearly stated otherwise), where we selected one class with clearly improved resolution for the VIPR complex, leaving 277,368 particles. These particles were further sorted with one round of 2D classification, leaving 208,701 particles. This particle stack was subjected to an additional round of 2D classification where we selected particles that showed clear secondary structure, giving 106,253 particles which were used for TOPAZ training (*18*). TOPAZ training and extraction yielded 72,047 particles, which were curated with one round of 2D classification giving 64,545 particles. These were pooled with a previously sorted stack of 208,701 particles and duplicates were removed, yielding 273,899 particles.

We used these particles for a one-class ab initio and non-uniform refinement. The map we generated from the non-uniform refinement was used for reference based motion correction before an additional non-uniform refinement using the motion-corrected particles, yielding a consensus map with an overall resolution of 2.99 Å. We generated a mask enclosing the five 5’-most subunits of this map, allowing additional space on the 5’ end of the VIPR complex. We used this mask for one round of 3D classification. Two classes had density for 11 subunits (113,361 particles), while two had density for 12 subunits (114,505 particles). Each subset was used for a final ab initio and non-uniform refinement, yielding two unique conformations: an 11-subunit class that refined to 3.17 Å overall resolution, and a 12-subunit class that refined to 3.13 Å overall resolution.

For SUSP1 VIPR incubated with an ssDNA target, 5,314 micrographs were motion-corrected with patch motion correction and contrast transfer function parameters were fit with patch CTF in cryoSPARC V4.6.2. 4,808,575 particles were selected by blob picking on micrographs CTF fit to 5.5 Å or better. One round of 2D classification yielded 1,858,232 particles which were used for a 3-class ab initio. One class with clear VIPR-like density was selected, giving 809,117 particles. This particle stack was further cleaned with a second three-class ab initio (maximum resolution of 3 Å, initial resolution of 10 Å, a Fourier step radius of 0.01, center structures in real space set to false, and a class similarity of 0. We note that these settings were used for all other ab initio jobs unless clearly stated otherwise). We selected one class with dramatically improved resolution, leaving 405,371 particles. We used this particle stack for a one-class ab initio followed by non-uniform refinement, which we used for reference-based motion correction, followed by an additional non-uniform refinement, yielding a consensus map with an overall resolution of 2.93 Å. Again, we generated a mask enclosing the five 5’-most subunits of this map, allowing additional space on the 5’ end of the VIPR complex. We used this mask for one round of 3D classification, requesting 4 separate classes. One class had clear density for 10 subunits (88,056 particles), two had clear density for 11 subunits (177,531 total particles), and the last class had clear density for 12 subunits (133,416 particles). For each conformational class, particles were pooled (if necessary), and used for one final ab initio and non-uniform refinement, yielding three unique conformations. The 10-subunit class refined to 3.08 Å overall resolution, the 11-subunit class refined to 3.0 Å overall resolution, and the 12-subunit refined to 3.03 Å overall resolution.

For SUSP1 VIPR incubated with an ssRNA target, 5,335 micrographs were motion-corrected with patch motion-correction and contrast transfer function parameters were fit with patch CTF in cryoSPARC v4.6.2. 5,000,764 particles were selected by blob picking on micrographs CTF fit to 6.0 Å or better. One round of 2D classification yielded 2,364,251 particles which were used for a 3-class ab initio. One class with clear VIPR-like density was selected with 1,095,595 particles, which we refer to as particle stack 1. The remaining particles were subject to one round of 2D classification per ab initio class yielding a separate stack of 1,006,283 particles which we refer to as particle stack 2.

Particle stack 1 was curated by two rounds of 2D classification, one three-class ab initio (maximum resolution of 3 Å, initial resolution of 10 Å, a Fourier step radius of 0.01, center structures in real space set to false, and a class similarity of 0. We note that these settings were used for all other ab initio jobs unless clearly stated otherwise), and one two class ab initio, yielding 226,589 particles. These particles were used for TOPAZ training and extraction, which yielded 516,726 particles. These particles were used for a two-class ab initio which yielded one high resolution class and one slightly lower resolution class. The particles from the higher resolution class were further curated with a two-class ab initio leaving 210,327 particles, while the particles in the lower resolution class were further curated by one round of 2D classification leaving 95,668 particles. These two sets of particles were pooled with the particles used for TOPAZ training and duplicates were removed with an interparticle distance of 100 Å, leaving 355,813 particles. These particles were polished using reference based motion correction in cryoSPARC v4.6.2. After reference based motion correction, one final two-class ab initio was run and 283,153 particles were selected for a final non uniform refinement yielding a closed VIPR conformation with an overall resolution of 2.76 Å.

Particle stack 2 was used for one three-class ab initio which resulted in one clear intermediate VIPR conformation (∼7-8 subunits) and two junk classes. A final non-uniform refinement yielded a map with an overall resolution of 2.94 Å. The resolution of this map was not improved with reference based motion correction.

For SUSP1 VIPR incubated with dsDNA, 9029 movies were motion-corrected with patch motion correction and contrast transfer function parameters were fit using patch CTF within cryoSPARC v4.6.2. 6,843,728 particles were selected using the blob picker on 4,217 micrographs CTF fit to 6.0 Å or better. One round of 2D classification yielded 3,006,924 particles, which were used for a 3-class ab initio. We selected one class with 1,217,801 particles that had clear VIPR-like density. Because this particle stack was so large, we elected to further clean this particle stack using a five-class ab initio (maximum resolution of 3 Å, initial resolution of 10 Å, a Fourier step radius of 0.01, center structures in real space set to false, and a class similarity of 0. We note that these settings were used for all other ab initio jobs unless clearly stated otherwise). We selected three classes with clear VIPR density, one class with 9 subunits comprising 303,503 particles, one class with 7 subunits comprising 294,707 particles, and a final class with 8 subunits comprising 339,842 particles.

Each of these particle stacks were used for separate 2-class ab initios. Of the resultant six ab initio classes, we selected five: A 9 subunit class comprising 162,512 particles, a 10 subunit class comprising 142,794 particles, a 7 subunit class, comprising 216,612 particles, a 8 subunit class comprising 203,735 particles, and a 8 subunit class comprising 135,098 particles. From each of these outputs, we ran a separate 3D classification job requesting 4 conformations using the mask from the parent non-uniform refinement. Our data suggested a wide range of potential intermediate classes with varying subunit counts, however at this stage, we still observed clear heterogeneity within classes, suggesting we needed an alternative method for filtering different subclasses. We grouped classes that had approximately 6-7 subunits (267,475 particles), 8-9 subunits (283,849), and 10-11 subunits (257,626 particles). We used each of these particle sets for separate 2-class ab initios. We still felt that class separation was not ideal. To combat this, we generated masks for each of the six resultant maps enclosing the five 5’-most subunits of the map, allowing additional space on the 5’ end of the VIPR complex. We used each of these masks as a focus mask for a separate 3D classification job requesting four conformations from each job, with the exception of one smaller class, where we requested three conformations. We regrouped the resultant 23 maps by number of clearly resolved subunits and used each pooled group for an ab initio, non-uniform refinement, reference based motion correction, and final non-uniform refinement. This approach gave us seven unique conformations, ranging from 5 subunits to 11 subunits, that resolves to 3.35 Å, 3.25 Å, 3.17 Å, 3.17 Å, 3.20 Å, 3.25 Å, and 3.53 Å for the 5-, 6-, 7-, 8-, 9-, 10-, and 11-subunit classes, respectively

For the *P. fulva* VIPR complex, movies were combined into one dataset of 7,483 micrographs and patch motion corrected in cryoSPARC. Contrast transfer function (CTF) parameters were fit with patch patch CTF within cryoSPARC. Micrographs were manually curated yielding 3,007 total micrographs. A circular blob pick was done with a diameter range of 60-250 Å yielding 6,911,653 initial particle picks that were curated by an inspect pick job to yield 2,576,174 particles. These were extracted at a 320-pixel box size for two-dimensional (2D) classification. Iterative rounds of 2D and 3D classification were conducted in cryoSPARC. and yielded a set of 480,707 particles. These particles were used to run a three-class ab initio (maximum resolution of 3 Å, initial resolution of 10 Å, a Fourier step radius of 0.01, center structures in real space set to false, and a class similarity of 0. We note that these settings were used for all other ab initio jobs unless clearly stated otherwise). Two structural classes appeared through this ab initio, an elongated filament class (129,804 particles, constricted conformation) and a truncated but stretched class (154,775, relaxed conformation). Each subset was subjected to one round of 2D classification before being used as templates for TOPAZ training and extraction. These two subsets were kept separate for the remainder of our processing of this dataset.

For the constricted conformation particles, we trained a TOPAZ model and extracted an additional 111,523 particles. These particles were pooled and duplicates were removed with an inter-particle distance of 100 Å. We investigated hunting for additional top views of the complex which appeared present within this dataset (data not shown). From our initial 480,707 particles, we selected all 2D classes that appeared to be top views and trained a TOPAZ model to drive picking on this conformation. This TOPAZ model yielded 1,345,948 particles, which were filtered through two rounds of 2D classification to yield 135,160 particles before re-training of TOPAZ with this new particle stack. This improved TOPAZ model yielded an additional stack of 138,384 particles. Each of these two particle stacks (135,160; TOPAZ model 1 and 138,384; TOPAZ model 2) were pooled with the particle stack above, and duplicates were removed with an inter-particle distance of 100 Å, yielding 397,749 particles. This particle stack was used for a 3-class ab initio, which generated two viable classes: an improved constricted conformation class and another relaxed conformation class. Non-uniform refinement and reference based motion correction of the constricted conformation resulted in a consensus constricted conformation that resolved to an overall resolution of 2.46 Å with C1 symmetry that was further improved by helical refinement to 2.30 Å. Helical refinement resulted in a slight loss in resolution at the nucleic acid level. This map was used as a reference for modeling of Vipr protomers, while the non-helically refined map was used as the primary refinement map. We observed pronounced density for a triplex within this construct. In an effort to improve this resolution, we generated a focus mask that targeted the center of the VIPR filament and used it for 3D classification. This gave us four classes with strong triplex density comprising 11 subunits (103,775 total particles) and one with weak triplex density comprising 10 subunits (26,173 total particles). Each of these particle stacks were used for a final ab initio and non-uniform refinement. The strong triplex particle stack resulted in a map that resolved to an overall resolution of 2.46 Å, while the weak triplex particle stack resulted in a map that resolved to an overall resolution of 2.70 Å.

For the relaxed conformation, all relaxed conformation particles were pooled and duplicates were removed with an interparticle diameter of 100 Å. Particles were sorted through two separate 2-class ab initios before a final non-uniform refinement, which resulted in a relaxed conformation that resolved to an overall resolution of 2.75 Å.

### Structure Modeling and Refinement

For the SUSP1 binary complex, an initial apo model was generated using AlphaFold3 prediction and multiple copies were rigid body fit to the density in Phenix (*19*). Model building was performed iteratively with manual adjustment in Coot (*20*), global real space refinement in Phenix (*21*), and geometry assessment in Molprobity (*22*). Target and vrRNA sequences were modeled *de novo* in Coot and ISOLDE.

SUSP1 ternary structures were built using the SUSP1 binary subunits as a starting point. Individual subunit models were docked using ChimeraX. Model building was performed iteratively with manual adjustment in Coot, global real space refinement in Phenix, and geometry assessment in Molprobity. Target and vrRNA sequences were modeled *de novo* in Coot and ISOLDE. The density for the third strand in models determined with ssDNA appeared markedly variable relative to that of the pre-unwound substrates, which precluded us from determining the exact base identity for this segment. As such, the third strand was modeled using the non-target strand from the previously modeled pre-unwound structures.

For the *P. fulva* complexes, an initial apo model was generated using AlphaFold3 prediction and multiple copies were rigid body fit to the density in Phenix. Model building was performed iteratively with manual adjustment in Coot, global real space refinement in Phenix, and geometry assessment in Molprobity. Target and vrRNA sequences were modeled *de novo* in Coot and ISOLDE. As the target substrate was a complement of endogenous targets (data not shown) and due to the lack of a fiducial marker like the 3’-pseudoknot seen in the SUSP1 VIPR complexes, we were unable to determine the exact register of the nucleic acids within the *P. fulva* VIPR structures. As such, non-conserved domains were modeled with the following logic: vrRNA NN domains were modeled as adenosines or cytosines, the target strand was modeled as complementary to the vrRNA with x_3_ bases being treated as cytosines or guanines as density permitted. The non-target strand was modeled largely as thymines, with intermittent guanines and cytosines as density permitted.

**Fig. S1.**
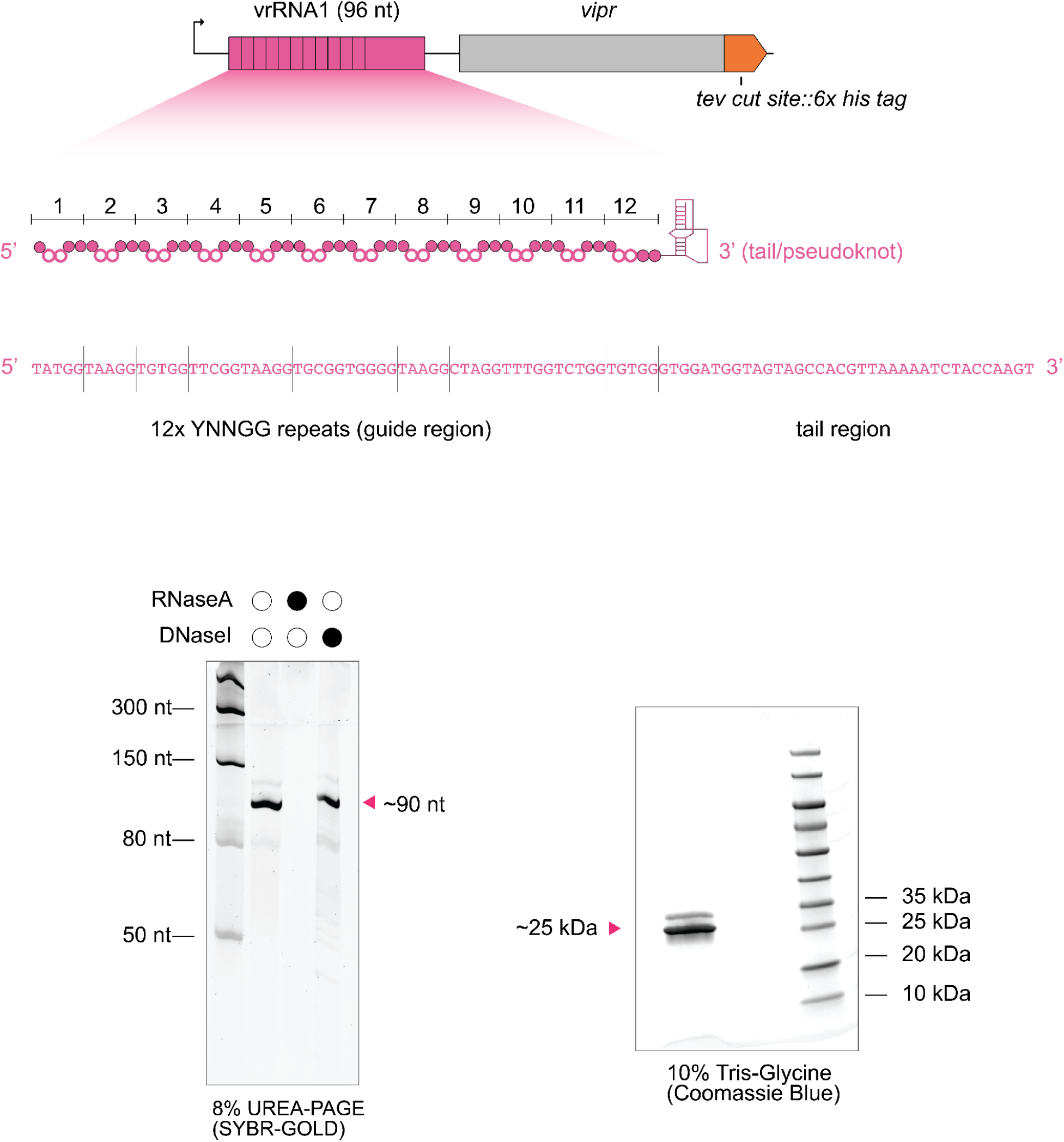
*E. coli* plasmid based heterologous expression and purification of SUSP1 VIPR.

**Fig. S2.**
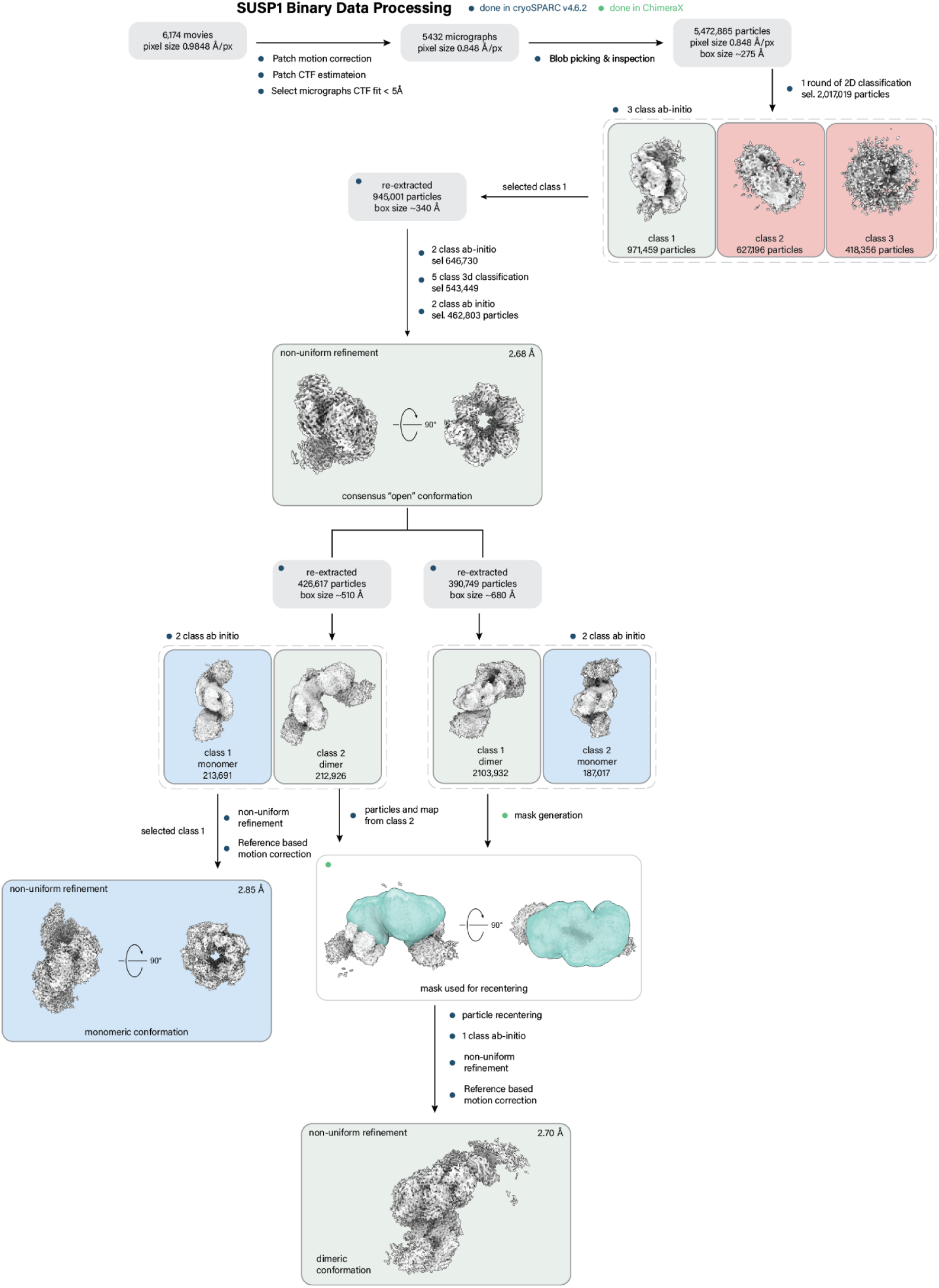
SUSP1 binary cryo-EM data processing.

**Fig. S3.**
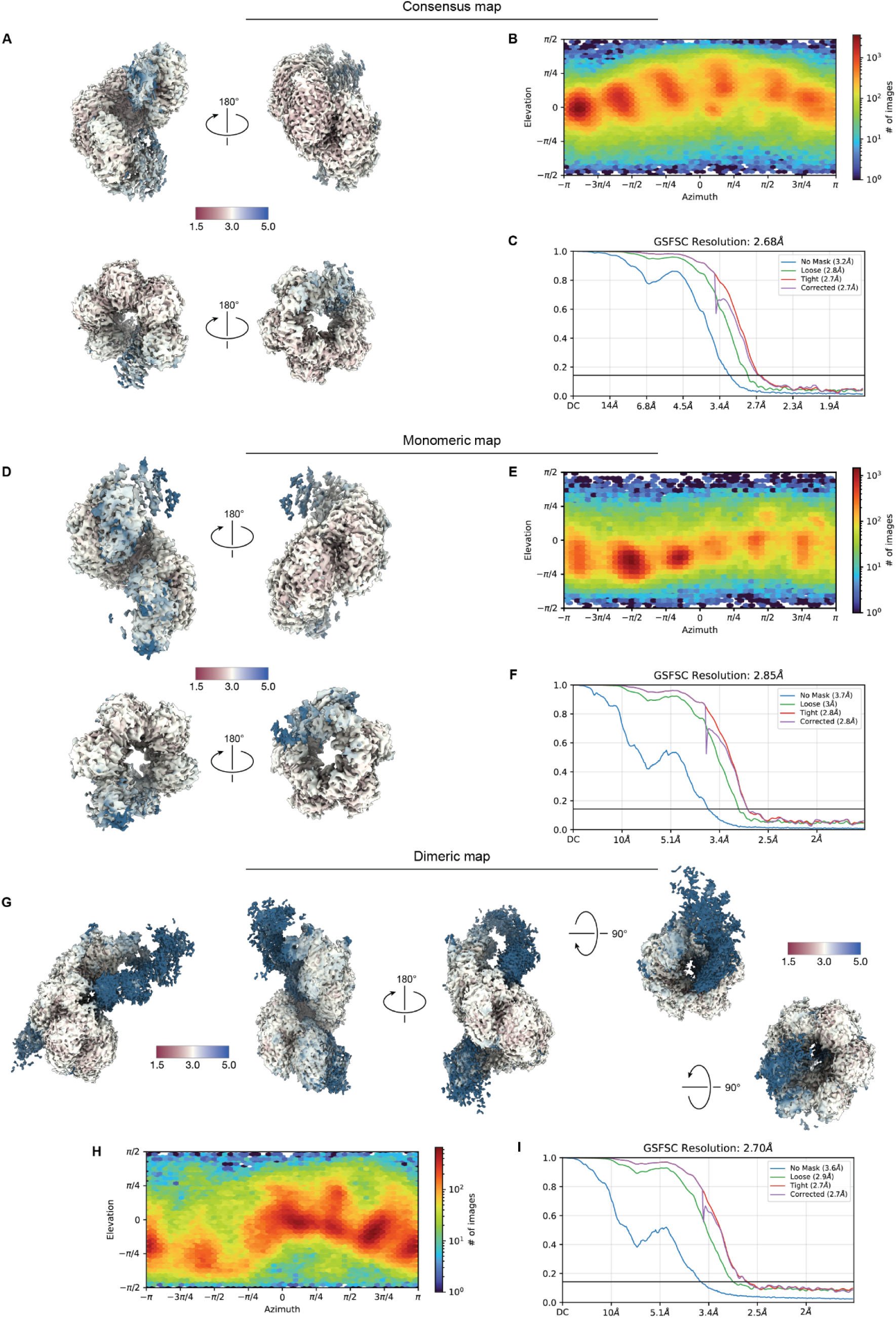
SUSP1 binary cryo-EM data validation. **(A)** Final sharpened consensus map colored by local resolution and viewed from the either side of the filament and from both the 3’ and 5’-ends. **(B)** View angle distribution of final particle stack. **(C)** Fourier shell correlation (FSC) versus resolution between half maps from final refinement for the consensus SUSP1 binary map. **(D)** Final sharpened monomeric map colored by local resolution and viewed from the either side of the filament and from both the 3’ and 5’-ends. **(E)** View angle distribution of final particle stack. **(F)** Fourier shell correlation (FSC) versus resolution between half maps from final refinement for the monomeric SUSP1 binary map. **(G)** Final sharpened dimeric map colored by local resolution and viewed from the either side of the filament and from both the 3’ and 5’-ends. **(H)** View angle distribution of final particle stack. **(I)** Fourier shell correlation (FSC) versus resolution between half maps from final refinement for the dimeric SUSP1 binary map.

**Fig. S4.**
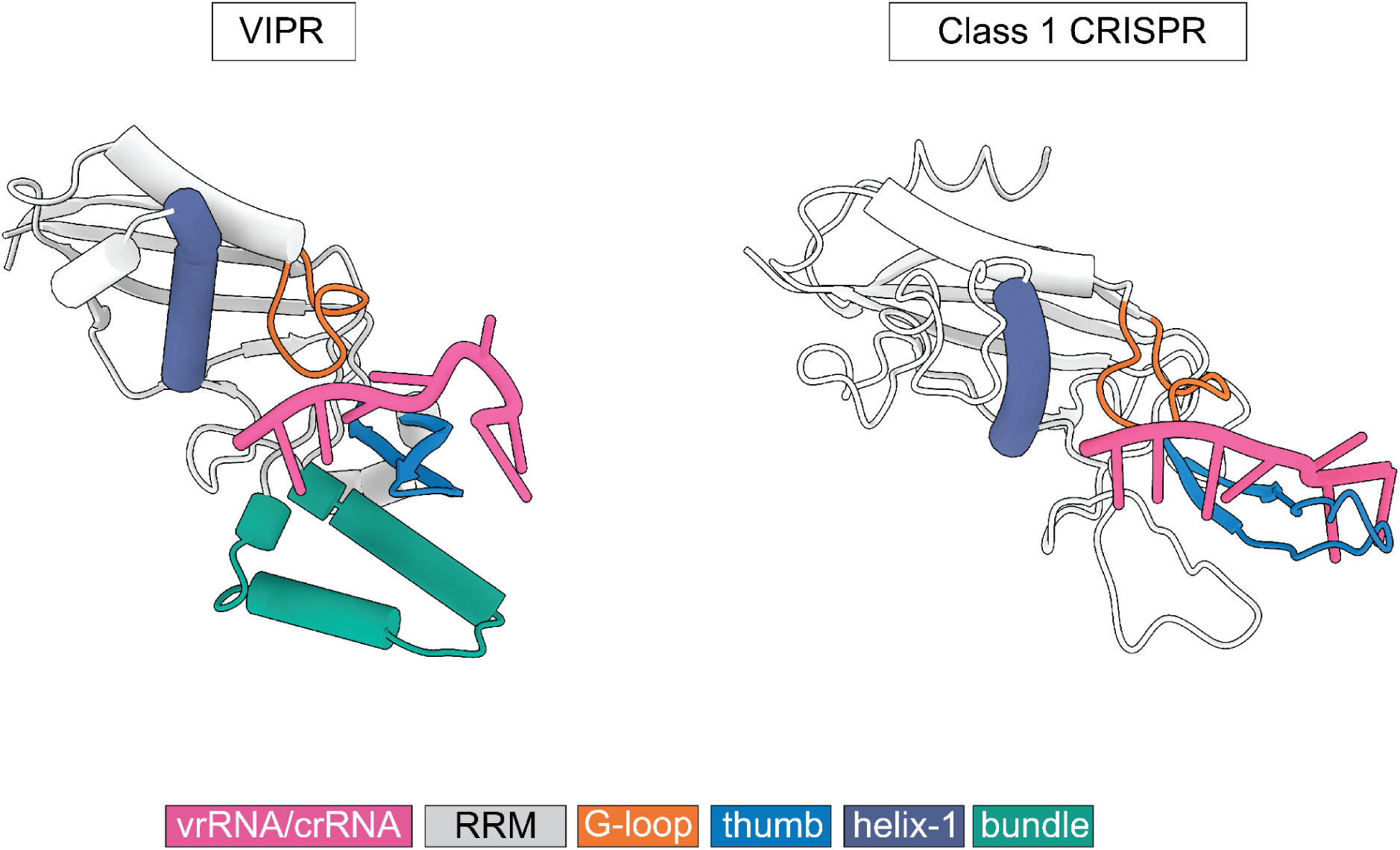
Comparison of guide RNA interaction between VIPR and a Class 1 CRISPR system. Class 1 CRISPR comparison is from type III-A, Cas7 subunit (Csm3), binding to its CRISPR RNA (crRNA) (PDBID: 6IFK).

**Supplementary Figure S5.**
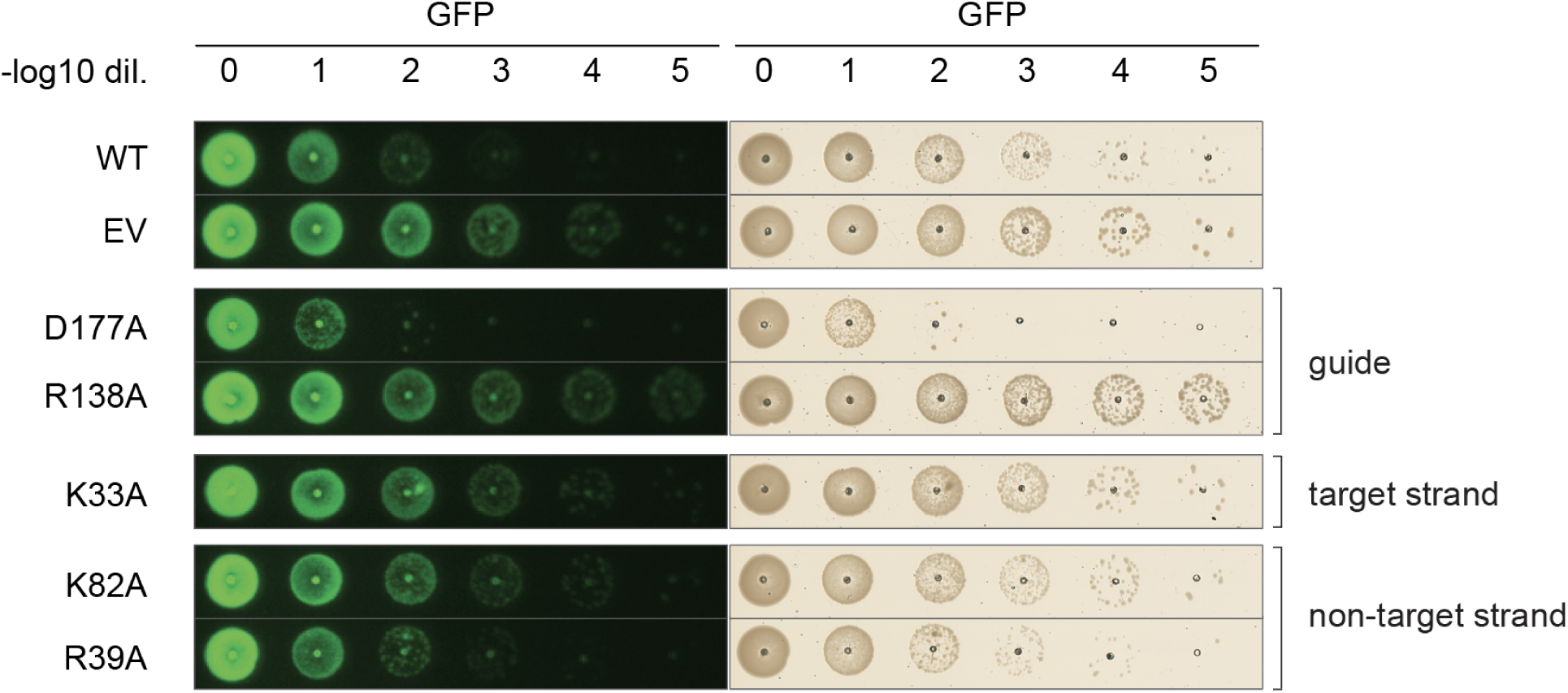
GFP repression assays comparing different mutations. Plasmid encoding VIPR (vrRNA and Vipr) were co-transformed with a GFP reporter plasmid with a VIPR target site downstream of the promoter to monitor transcriptional repression. The blue light channel is shown on the left to show GFP expression levels, and the brightfield channel is shown on the right to show cell density.

**Fig. S6.**
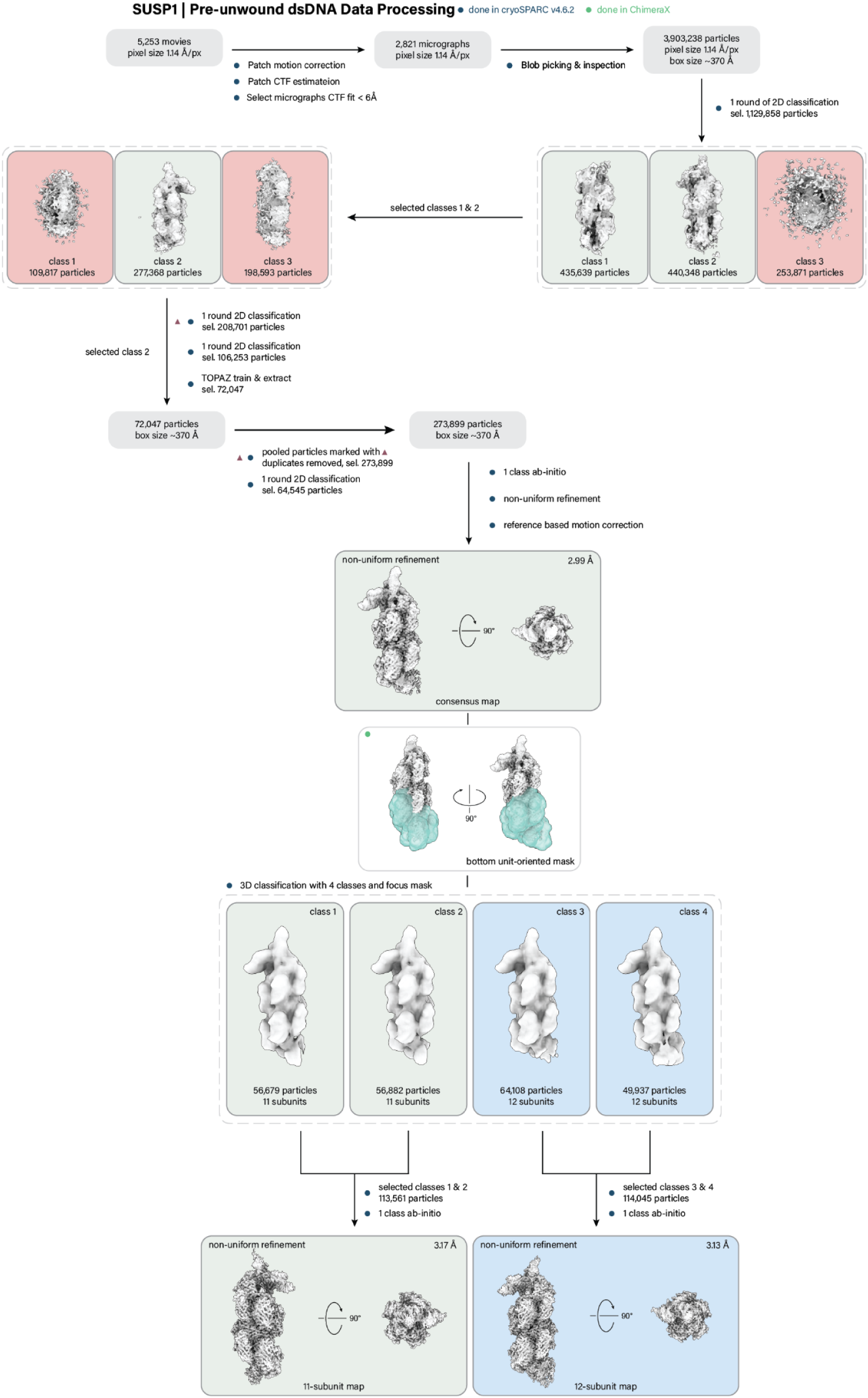
SUSP1 VIPR with pre-unwound (unpaired) dsDNA ternary cryo-EM data processing.

**Fig S7.**
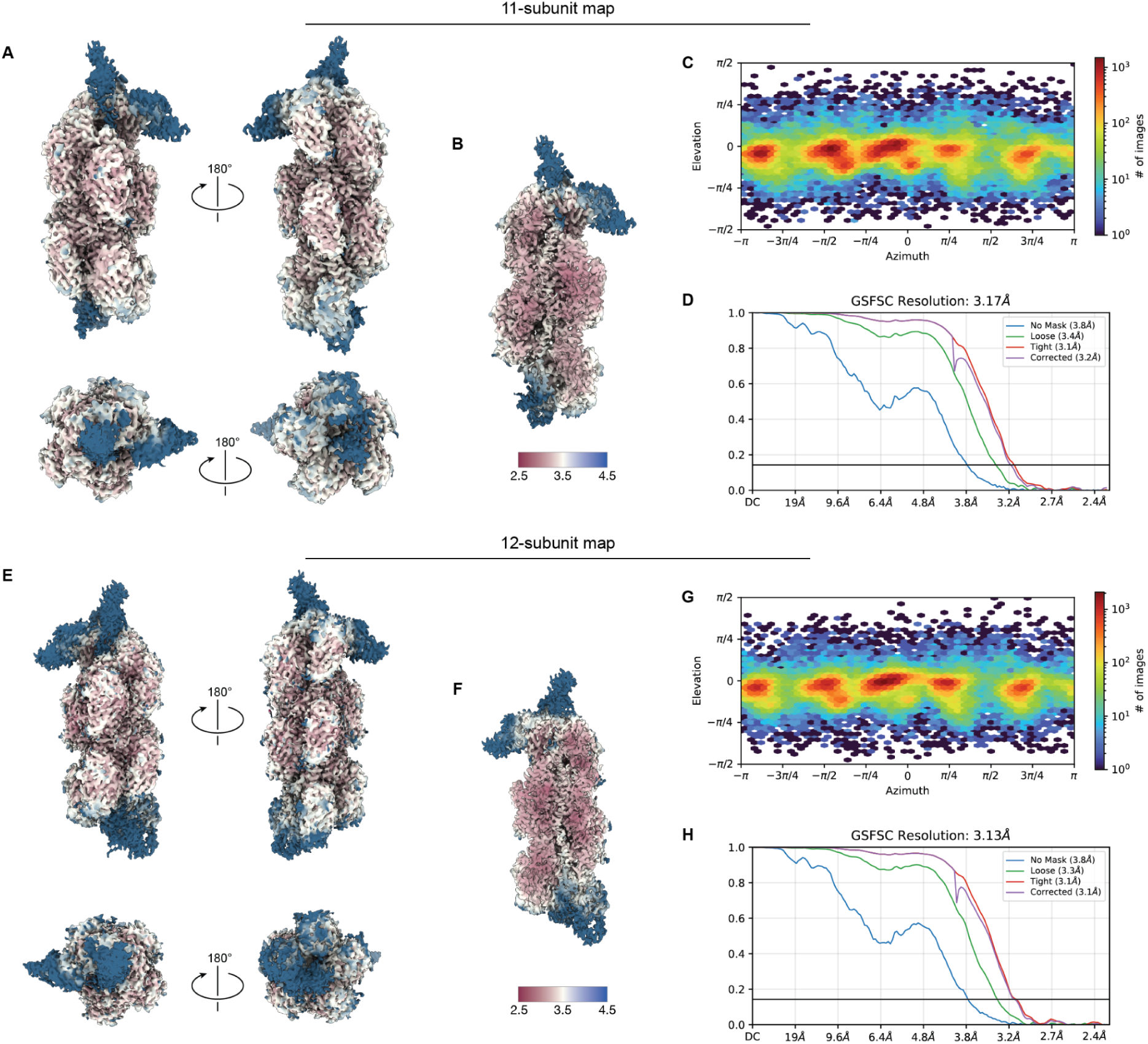
SUSP1 VIPR with pre-unwound (unpaired) dsDNA ternary cryo-EM data validation. **(A)** Final sharpened 11-subunit map colored by local resolution and viewed from the either side of the filament and from both the 3’ and 5’-ends. **(B)** Cutaway map showing resolution within the filament cavity of the 11-subunit map. **(C)** View angle distribution of final particle stack. **(D)** Fourier shell correlation (FSC) versus resolution between half maps from final refinement for the 11-subunit map. **(E)** Final sharpened 12-subunit map colored by local resolution and viewed from the either side of the filament and from both the 3’ and 5’-ends. **(F)** Cutaway map showing resolution within the filament cavity of the 12-subunit map. **(G)** View angle distribution of final particle stack. **(H)** Fourier shell correlation (FSC) versus resolution between half maps from final refinement for the 12-subunit map.

**Fig. S8.**
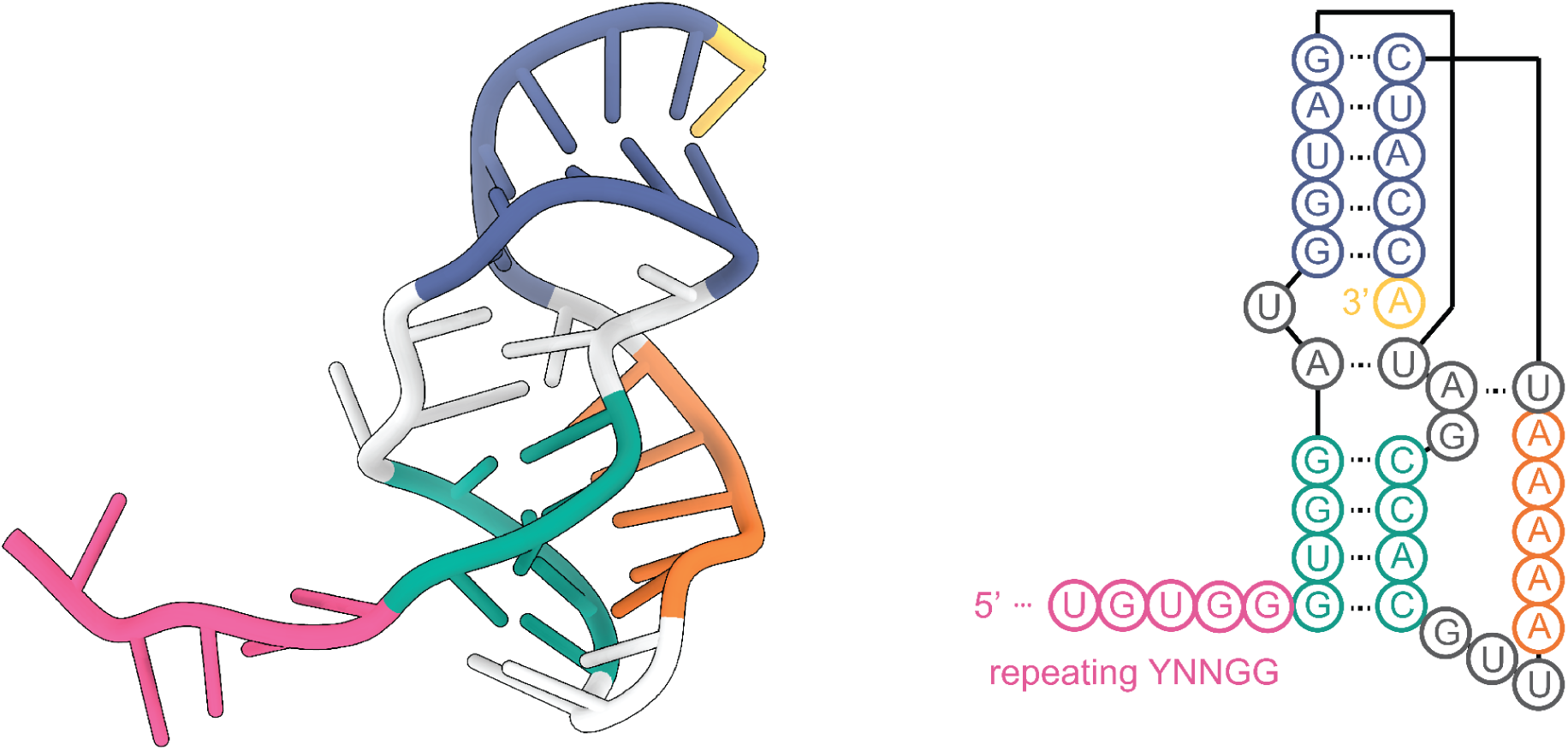
Molecular structure of SUSP1 vrRNA 3’ pseudoknot.

**Fig. S9.**
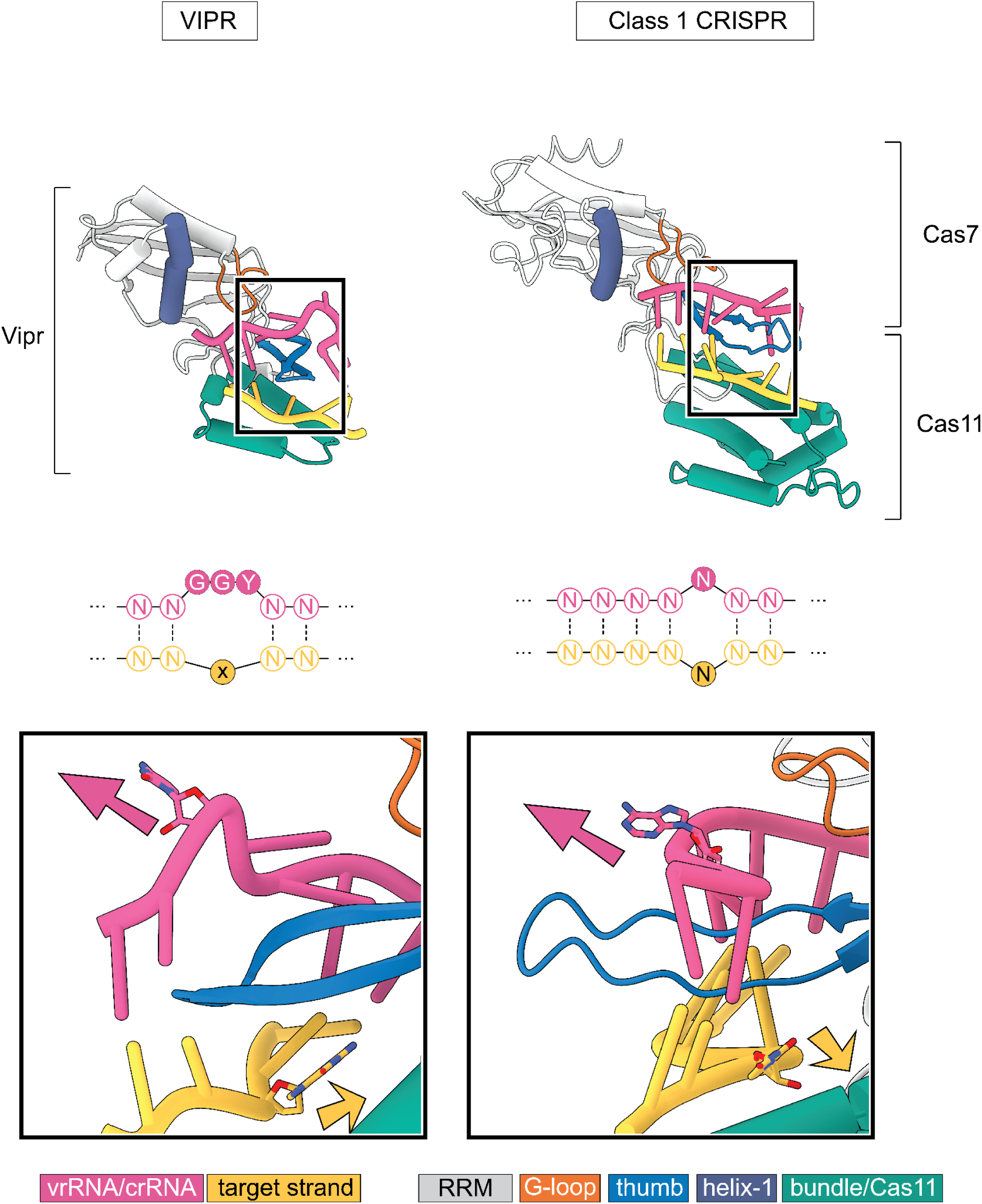
Comparison of target engagement in VIPR and a Class 1 CRISPR system. Class 1 CRISPR comparison is from type III-A, Cas7 subunit (Csm3), binding to its crRNA and target RNA (PDBID: 6IFK).

**Fig. S10.**
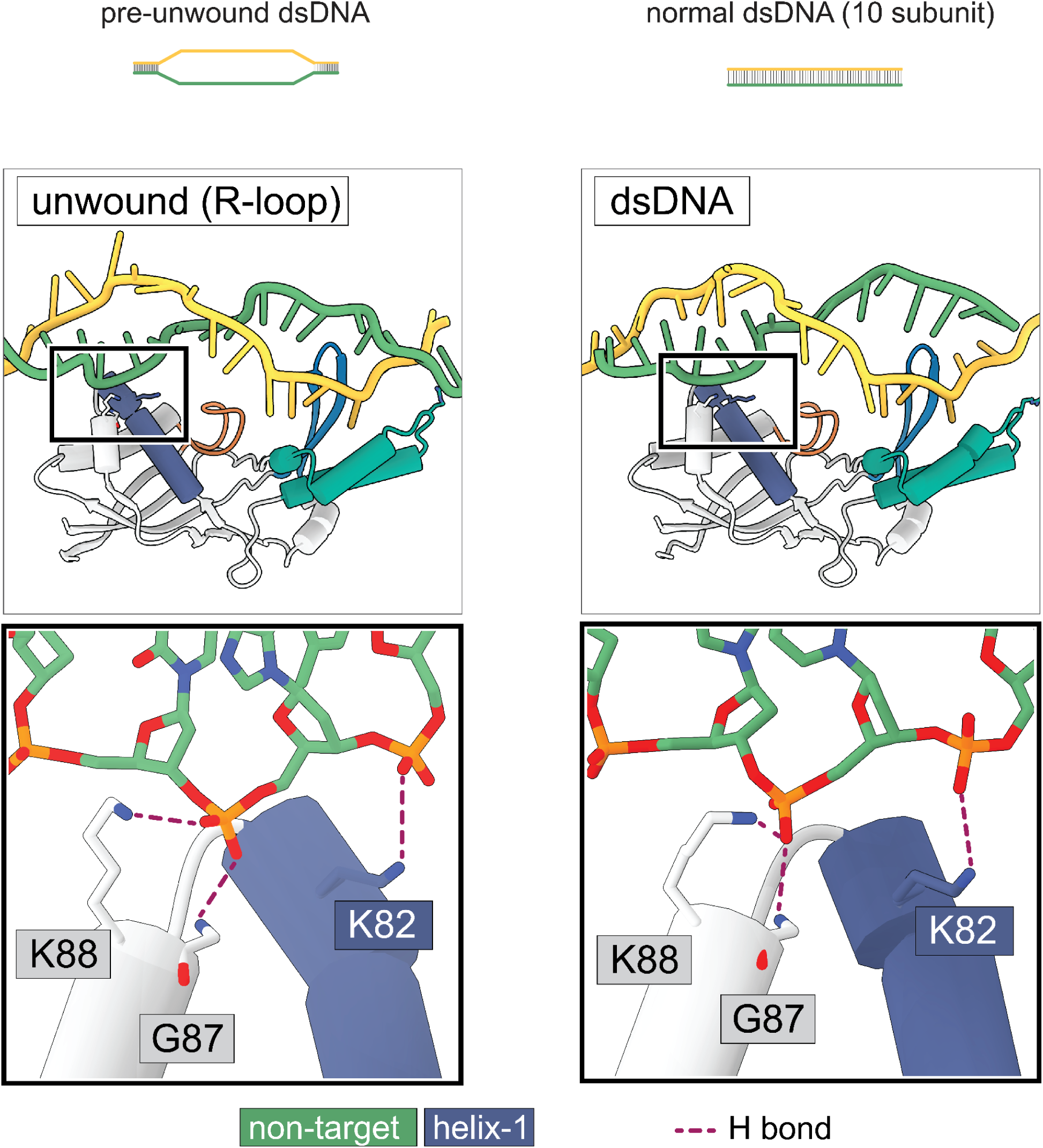
Non-target strand interaction in pre-unwound DNA and normal dsDNA formed ternary complex.

**Fig. S11.**
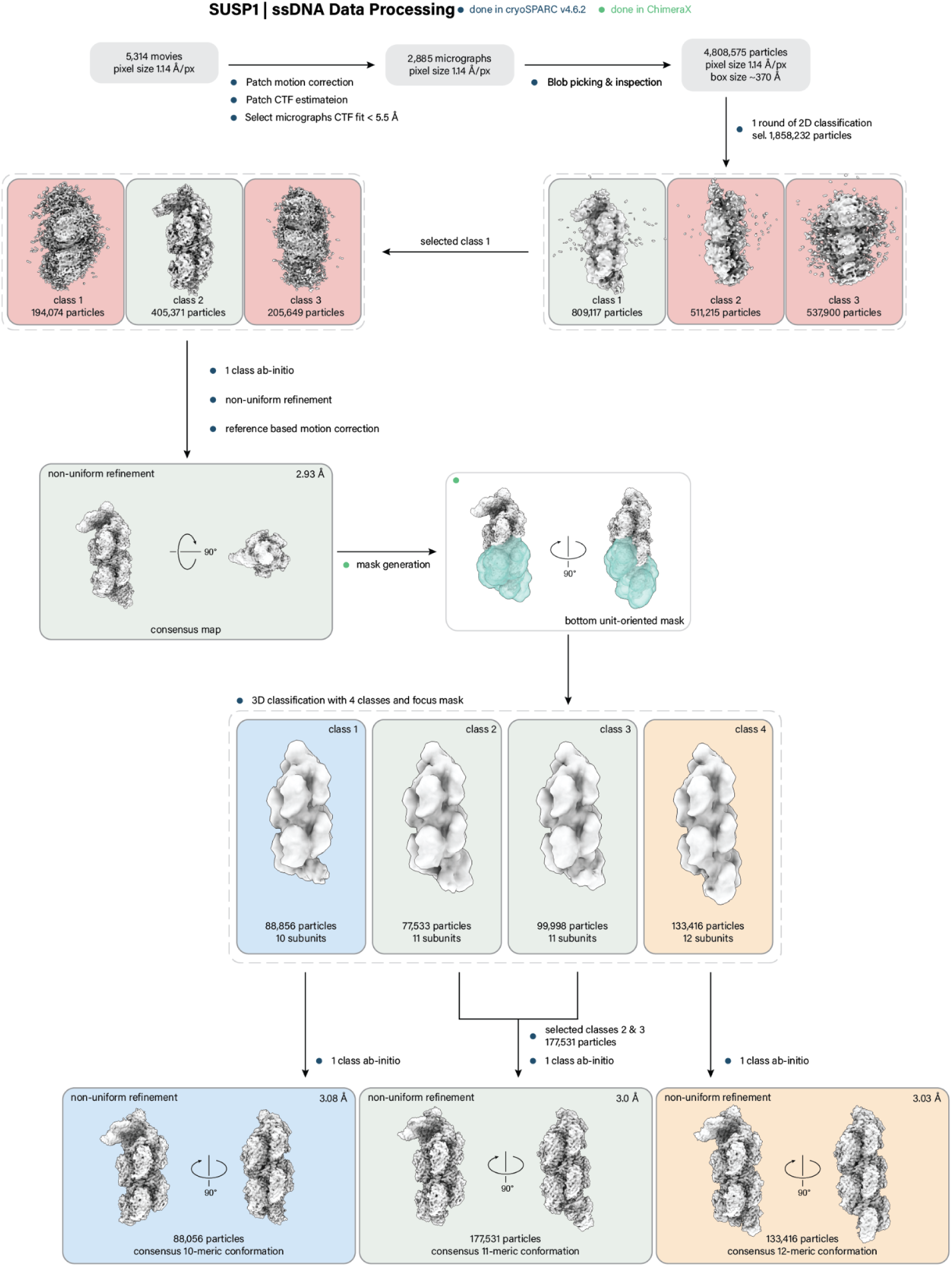
SUSP1 ssDNA ternary cryo-EM data processing.

**Fig. S12.**
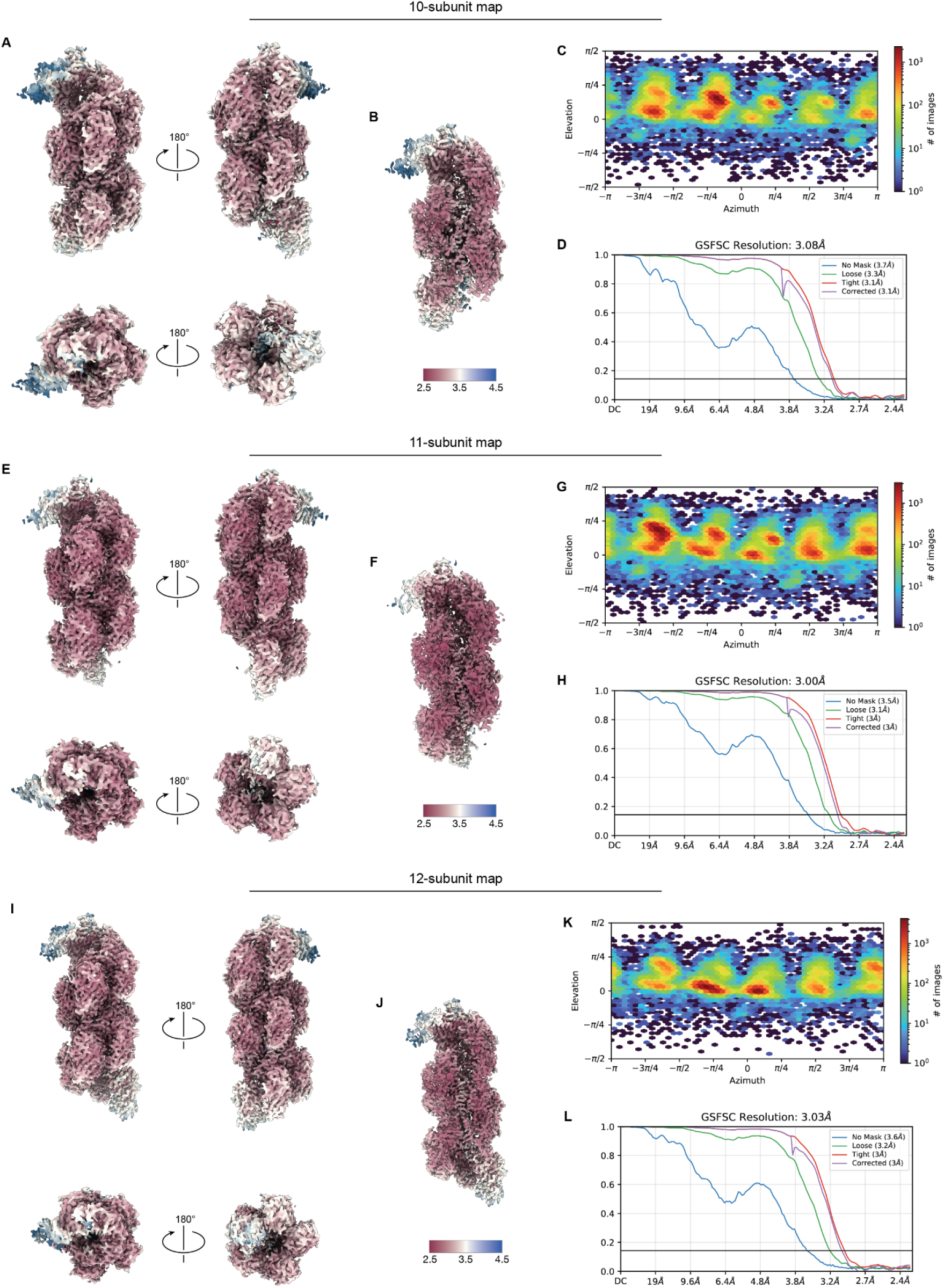
SUSP1 ssDNA ternary cryo-EM data validation. **(A)** Final sharpened 10-subunit map colored by local resolution and viewed from the either side of the filament and from both the 3’ and 5’-ends. **(B)** Cutaway map showing resolution within the filament cavity of the 10-subunit map. **(C)** View angle distribution of final particle stack. **(D)** Fourier shell correlation (FSC) versus resolution between half maps from final refinement for the 10-subunit map. **(E)** Final sharpened 11-subunit map colored by local resolution and viewed from the either side of the filament and from both the 3’ and 5’-ends. **(F)** Cutaway map showing resolution within the filament cavity of the 11-subunit map. **(G)** View angle distribution of final particle stack. **(H)** Fourier shell correlation (FSC) versus resolution between half maps from final refinement for the 11-subunit map. **(I)** Final sharpened 12-subunit map colored by local resolution and viewed from the either side of the filament and from both the 3’ and 5’-ends. **(J)** Cutaway map showing resolution within the filament cavity of the 12-subunit map. **(K)** View angle distribution of final particle stack. **(L)** Fourier shell correlation (FSC) versus resolution between half maps from final refinement for the 12-subunit map.

**Fig. S13.**
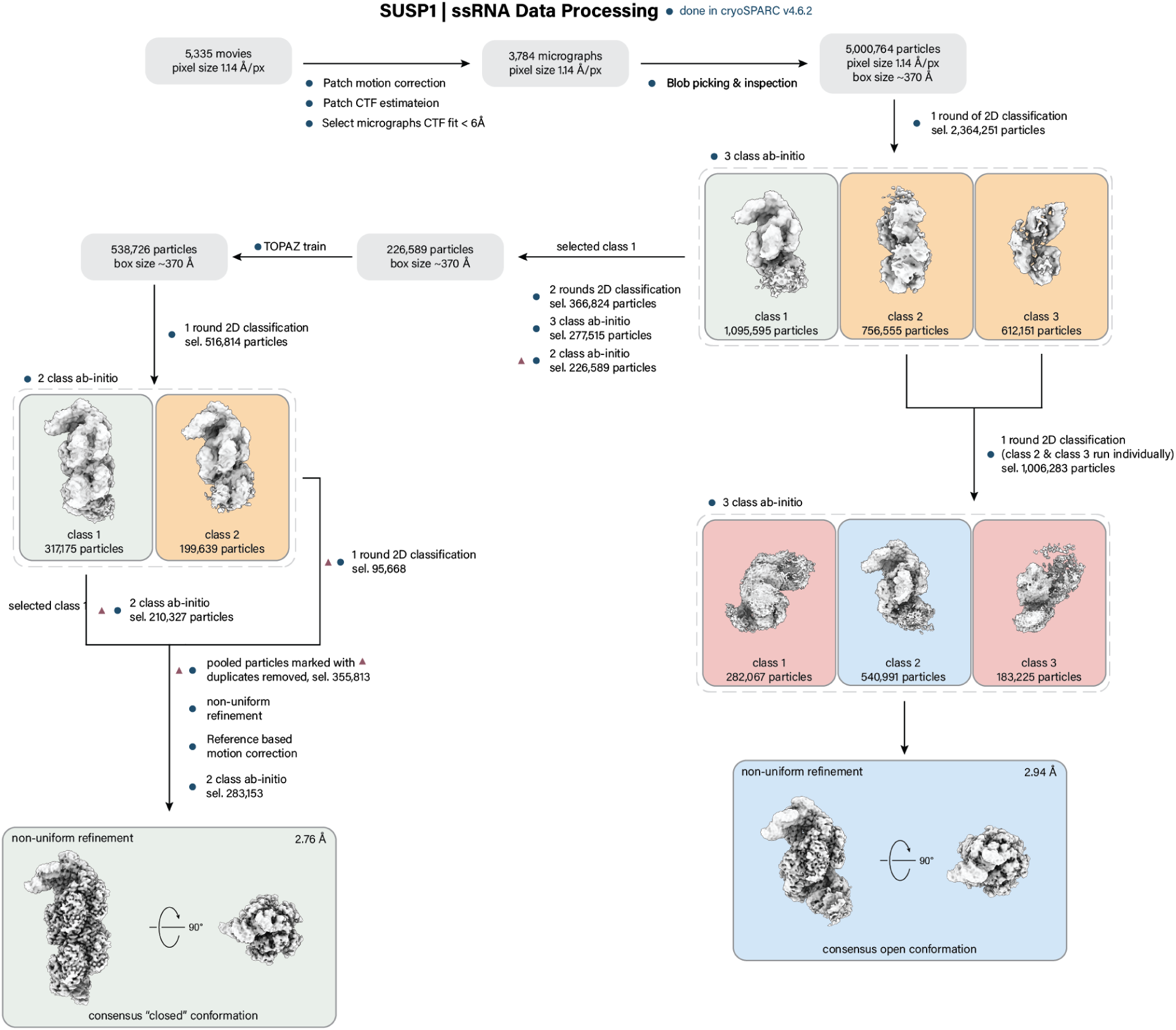
SUSP1 ssRNA ternary cryo-EM data processing.

**Fig. S14.**
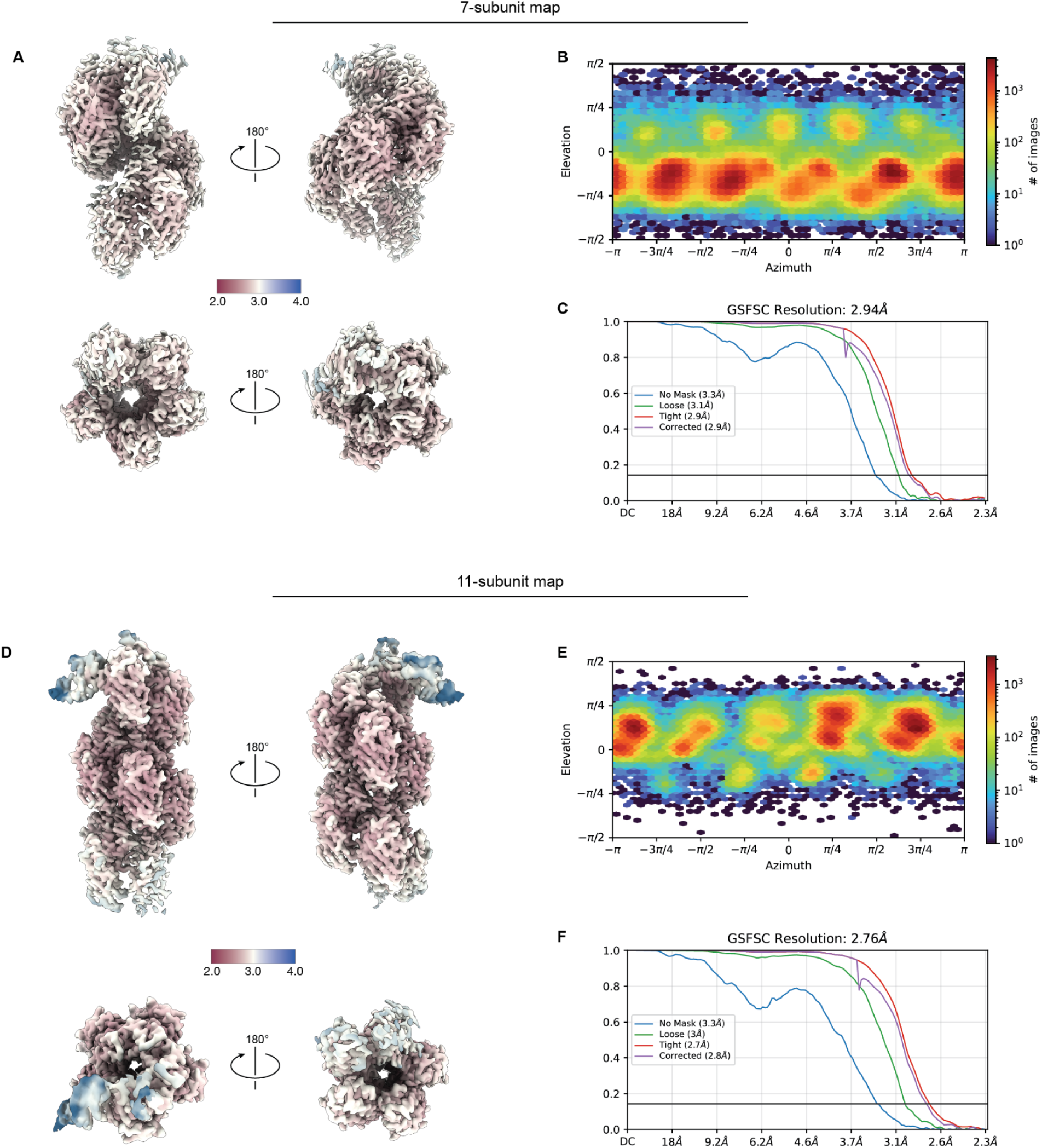
SUSP1 ssRNA ternary cryo-EM data validation. **(A)** Final sharpened 7-subunit map colored by local resolution and viewed from the either side of the filament and from both the 3’ and 5’-ends. **(B)** View angle distribution of final particle stack. **(C)** Fourier shell correlation (FSC) versus resolution between half maps from final refinement for the 7-subunit map. **(D)** Final sharpened 11-subunit map colored by local resolution and viewed from the either side of the filament and from both the 3’ and 5’-ends. **(E)** View angle distribution of final particle stack. **(F)** Fourier shell correlation (FSC) versus resolution between half maps from final refinement for the 11-subunit map.

**Fig. S15.**
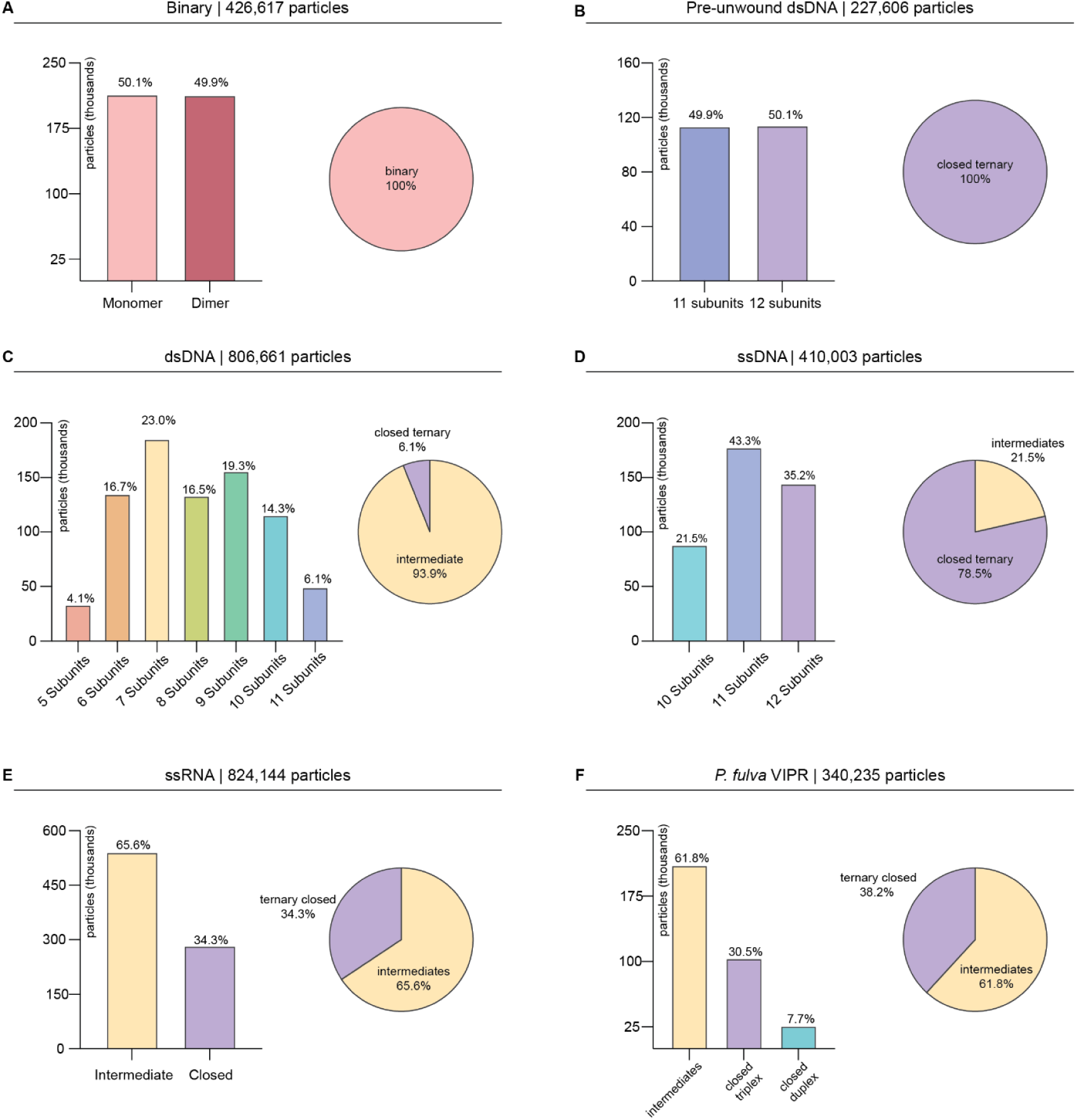
Particle distributions by construct. Final particle distributions for refined particle stacks. Left, bar graph showing particle count and percent of total particles for each sub-state within each dataset. Right, distribution of particles between binary (no target), intermediate ternary states (5-10 subunits), and closed ternary states (11-12 subunits). **(A**) Particle distribution SUSP1 binary (426,617 particles). **(B)** Particle distribution for SUSP1 incubated with a pre-unwound dsDNA target (227,606 particles). **(C)** Particle distribution for SUSP1 incubated with a dsDNA target (806,661 particles). **(D)** Particle distribution for SUSP1 incubated with an ssDNA target (410,003 particles). **(E)** Particle distribution for SUSP1 incubated with an ssRNA target (824,144 particles). **(F)** Particle distribution for *P. fulva* complexes (340,235 particles).

**Fig. S16.**
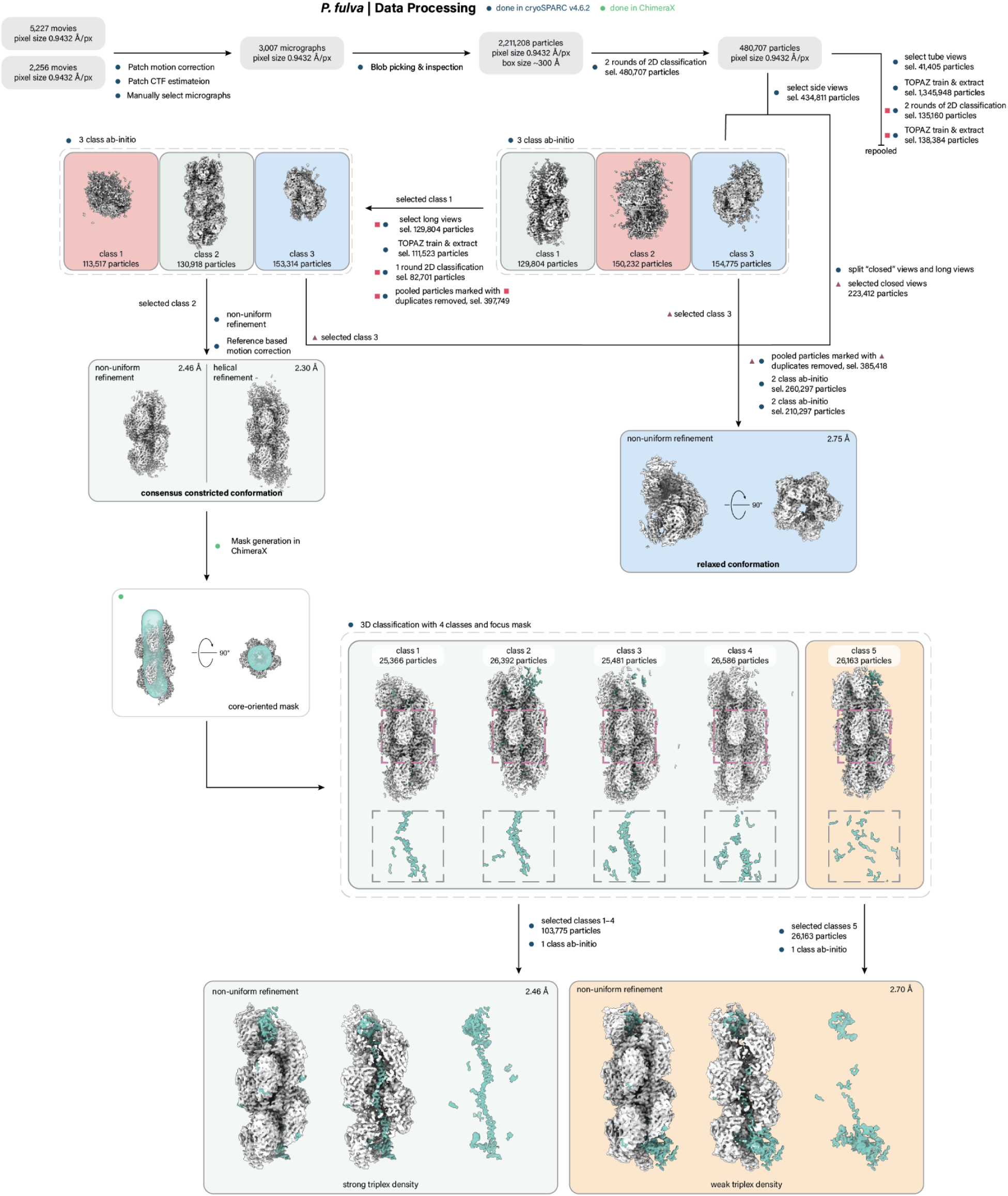
*P. fulva* ternary cryo-EM data processing.

**Fig. S17.**
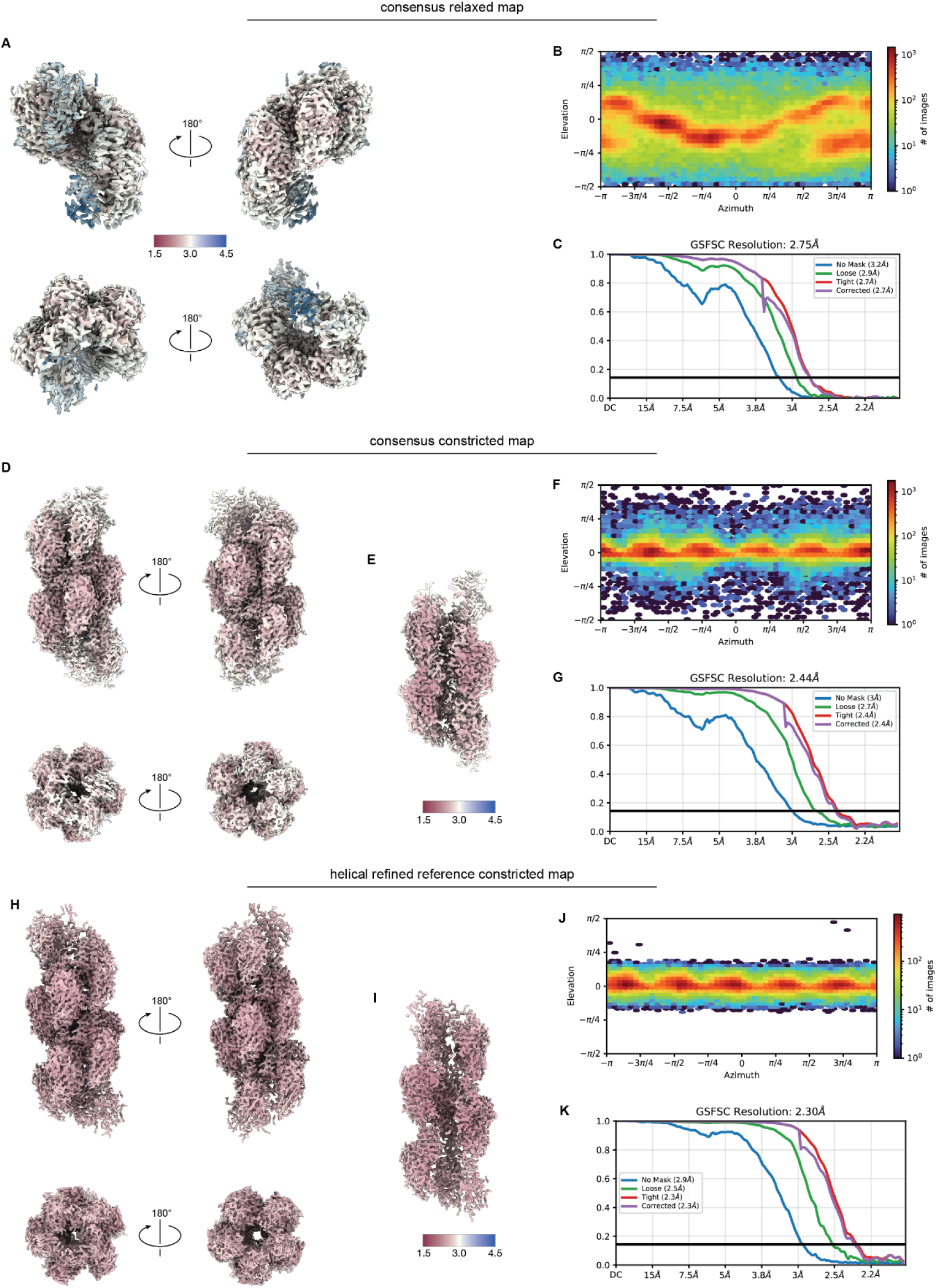
*P. fulva* ternary cryo-EM data validation part one. **(A)** Final sharpened relaxed conformation map colored by local resolution and viewed from the either side of the filament and from both the 3’ and 5’-ends. **(B)** View angle distribution of final particle stack. **(C)** Fourier shell correlation (FSC) versus resolution between half maps from final refinement for the relaxed conformation map. **(D)** Final sharpened consensus constricted conformation map colored by local resolution and viewed from the either side of the filament and from both the 3’ and 5’-ends. **(E)** Cutaway map showing resolution within the filament cavity of the consensus constricted conformation map. **(F)** View angle distribution of final particle stack. **(G)** Fourier shell correlation (FSC) versus resolution between half maps from final refinement for the consensus constricted conformation map. **(H)** Final sharpened helical-refined consensus constricted conformation reference map colored by local resolution and viewed from the either side of the filament and from both the 3’ and 5’-ends. **(I)** Cutaway map showing resolution within the filament cavity of the helical-refined consensus constricted conformation reference map. **(J)** View angle distribution of final particle stack. **(K)** Fourier shell correlation (FSC) versus resolution between half maps from final refinement for the helical-refined consensus constricted conformation reference map.

**Fig. S18.**
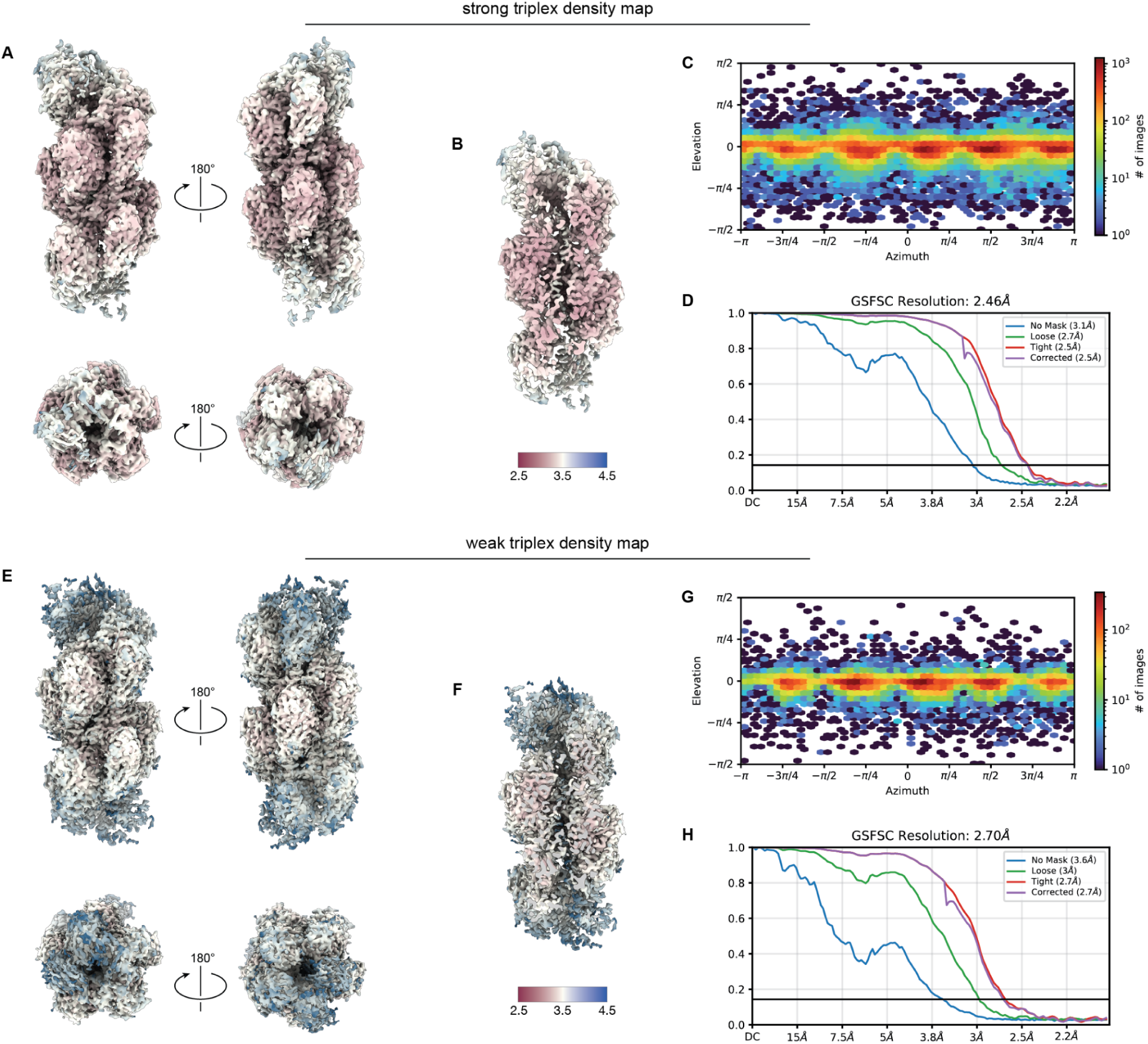
*P. fulva* ternary cryo-EM data validation part two. **(A)** Final sharpened strong-triplex map colored by local resolution and viewed from the either side of the filament and from both the 3’ and 5’-ends. **(B)** Cutaway map showing resolution within the filament cavity of the strong-triplex map. **(C)** View angle distribution of final particle stack. **(D)** Fourier shell correlation (FSC) versus resolution between half maps from final refinement for the strong-triplex map. **(E)** Final sharpened weak-triplex map colored by local resolution and viewed from the either side of the filament and from both the 3’ and 5’-ends. **(F)** Cutaway map showing resolution within the filament cavity of the weak-triplex map. **(G)** View angle distribution of final particle stack. **(H)** Fourier shell correlation (FSC) versus resolution between half maps from final refinement for the weak-triplex map.

**Fig. S19.**
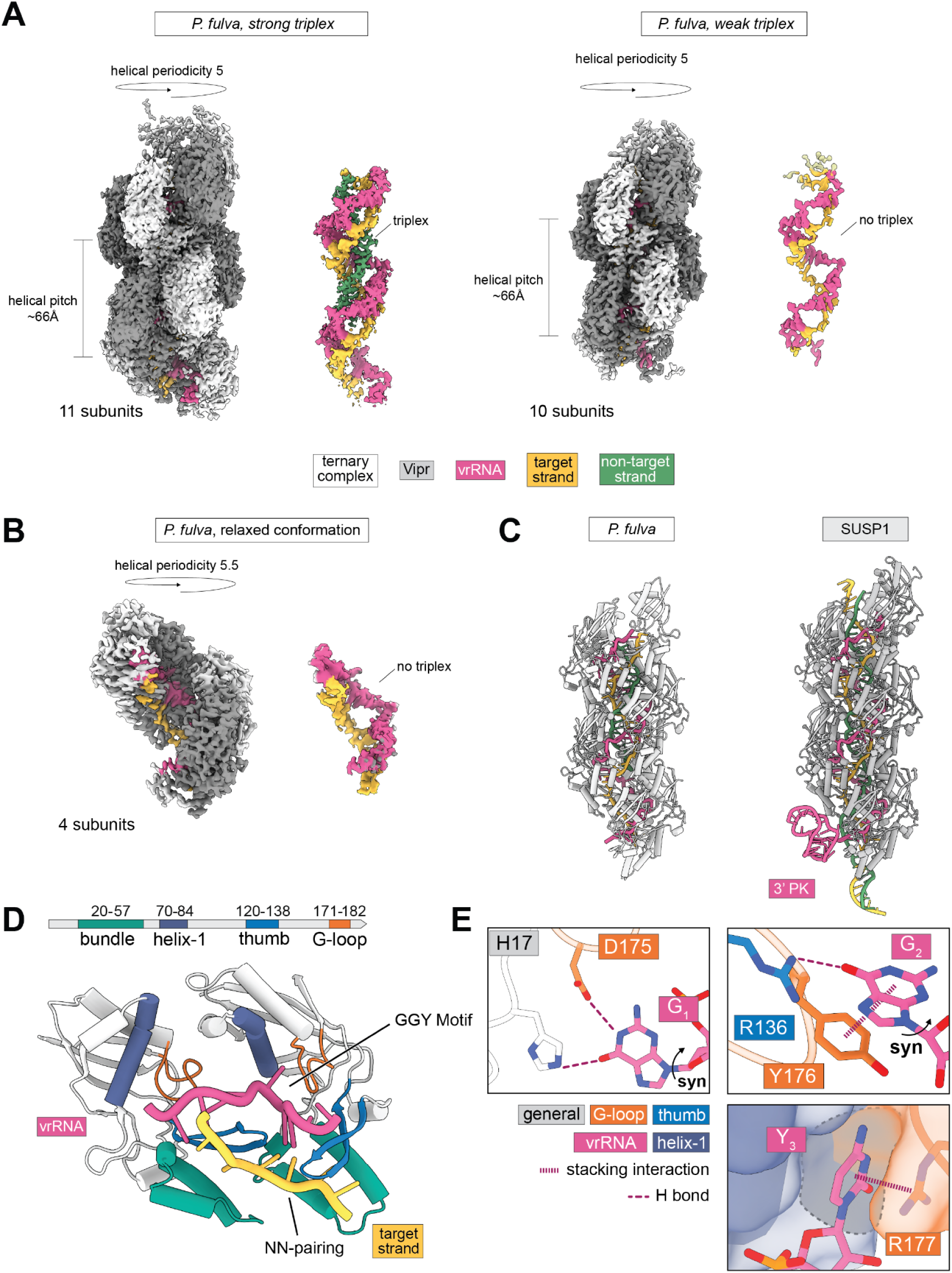
*P. fulva* VIPR forms an elongated superhelix with conserved vrRNA coordination and target engagement. **(A)** Cryo-EM map of two *P. fulva* VIPR ternary complexes, left depicting a class with strong triplex density and right depicting a class with weak triplex density. Loss of triplex density corresponds with a reduction in subunit count from 11 subunits to 10 subunits. **(B)** Cryo-EM map of a relaxed conformation of the *P. fulva* VIPR ternary complex with four subunits. **(C)** Comparison of the overall architecture of *P. fulva* VIPR (left) to SUSP1 VIPR (right). **(D)** *P. fulva* Vipr protomer domain architecture (top) and interactions with vrRNA and target DNA shown through two Vipr protomers spanning ∼2 vrRNA repeat units (bottom). **(E)** Base-specific contacts at G_1_, G_2_, and Y_3_ positions. Hydrogen bonds (H bond) and stacking interactions indicated. G_1_ is recognized by hydrogen bonds from the RRM β-sheet (H17) and the G-loop (D175), whereas G_2_ is cradled between the thumb (salt bridge, R136) and G-loop (π-stacking, Y176). Within the second protomer, Y_3_ is held in a narrow pocket formed by helix-1 (steric restriction, S75, A79) and the G-loop (cation-π, R177, steric restriction, C171, V173) that sterically excludes purines.

**Fig. S20.**
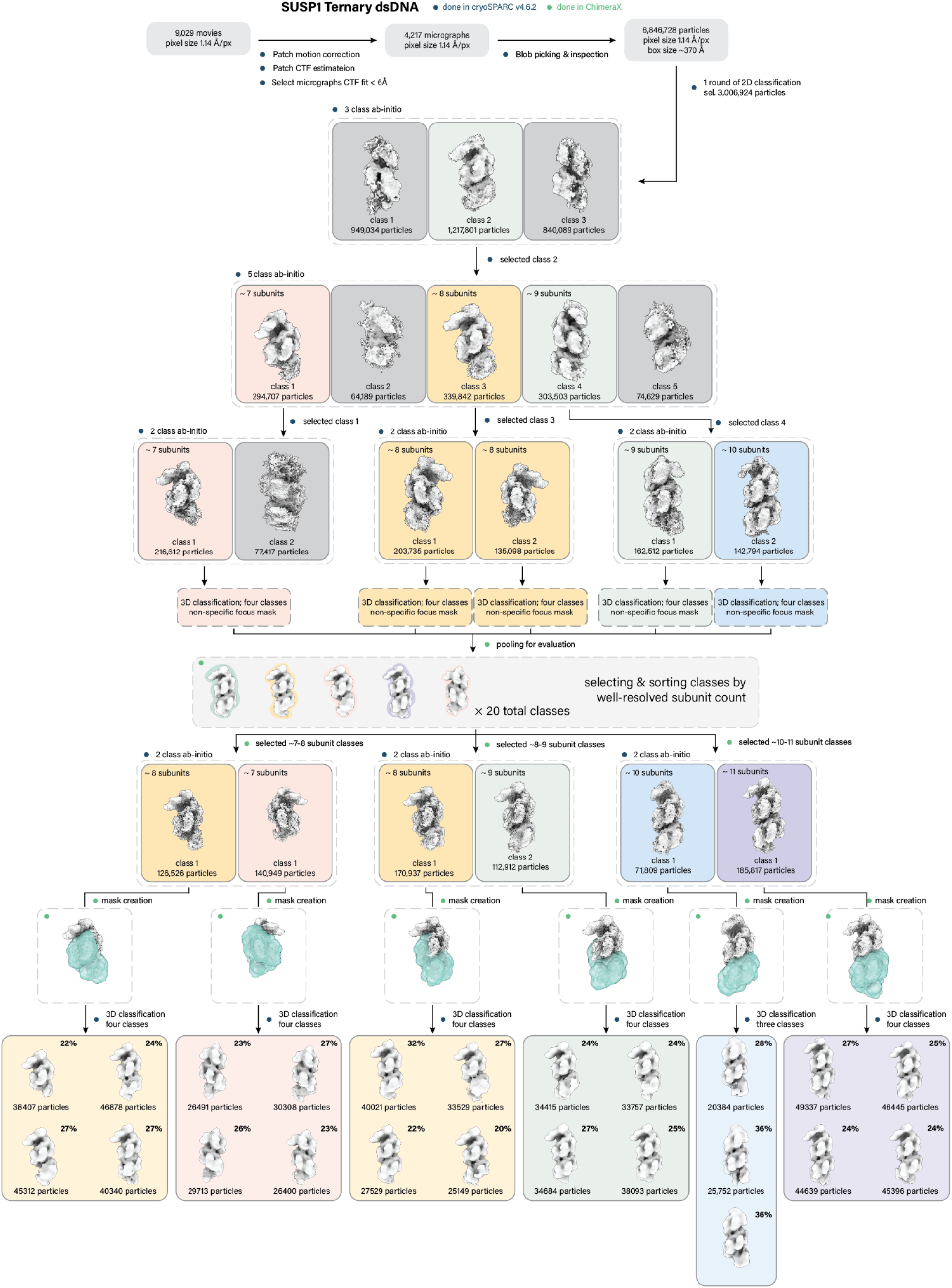
SUSP1 dsDNA ternary cryo-EM data processing part one. Data processing pipeline for SUSP1 incubated with a double-stranded DNA target, depicting processing from micrograph curation until final 3D classification step.

**Fig. S21.**
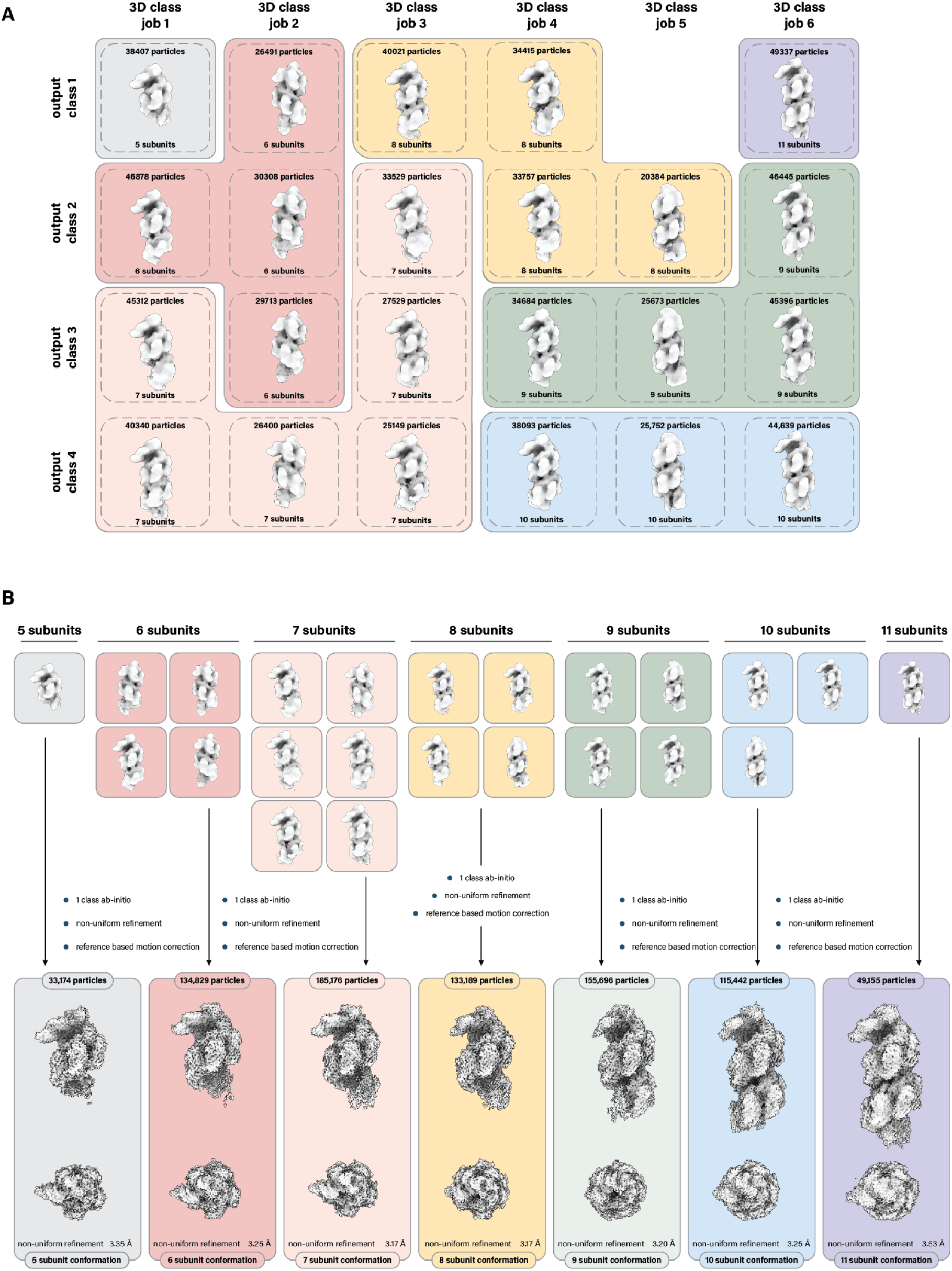
SUSP1 dsDNA ternary cryo-EM data processing part two. **(A)** Sorting of 3D classification outputs shown at the end of supplemental figure 17. Subunit count was calculated following a non-uniform refinement of the output class rather than the number of subunits present in the classification output. **(B)** Final processing steps for each separate class of particles.

**Fig. S22.**
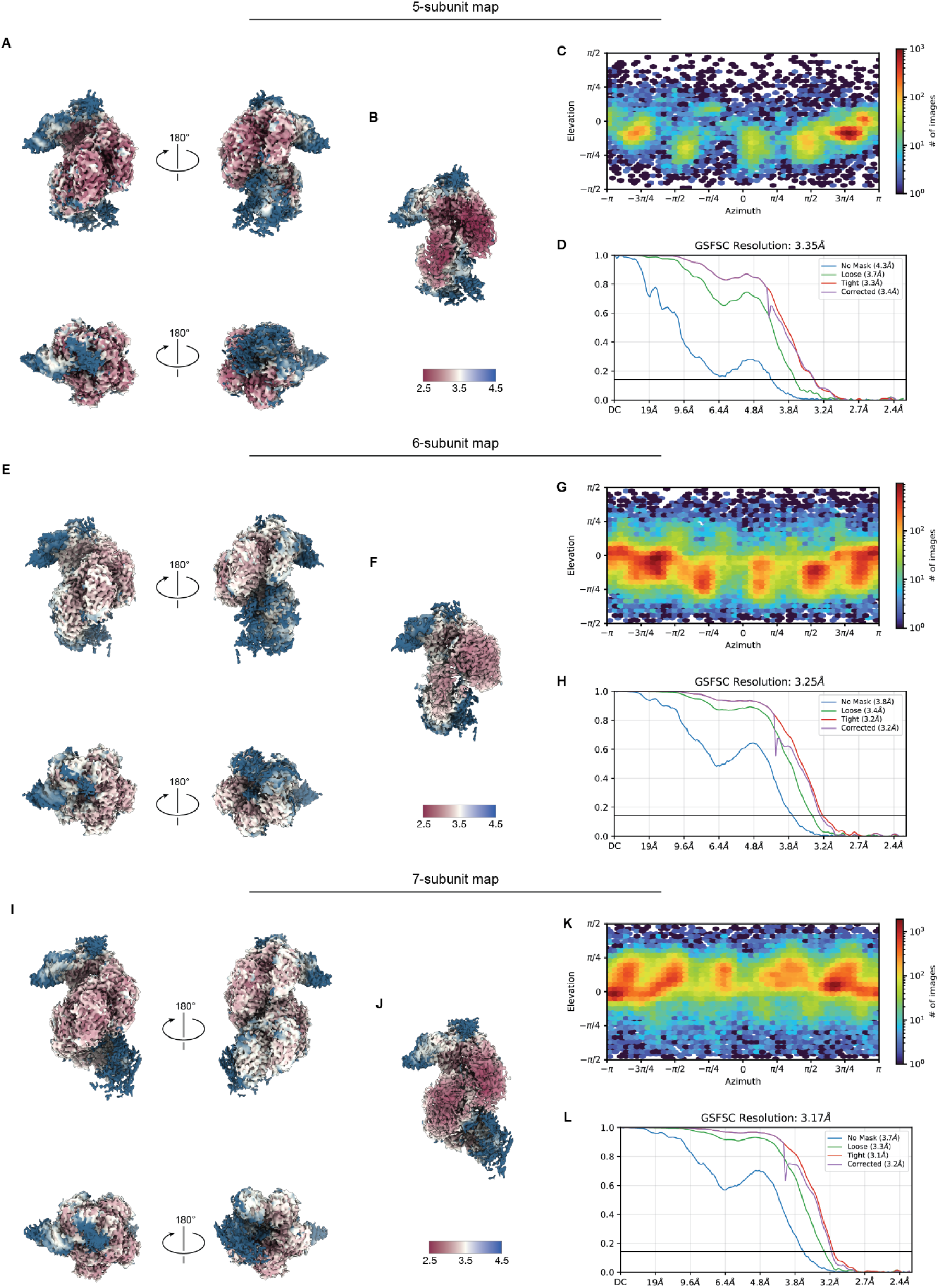
SUSP1 dsDNA ternary cryo-EM data validation for 5, 6, and 7 subunit conformations. **(A)** Final sharpened 5-subunit map colored by local resolution and viewed from the either side of the filament and from both the 3’ and 5’-ends. **(B)** Cutaway map showing resolution within the filament cavity of the 5-subunit map. **(C)** View angle distribution of final particle stack. **(D)** Fourier shell correlation (FSC) versus resolution between half maps from final refinement for the 5-subunit map. **(E)** Final sharpened 6-subunit map colored by local resolution and viewed from the either side of the filament and from both the 3’ and 5’-ends. **(F)** Cutaway map showing resolution within the filament cavity of the 6-subunit map. **(G)** View angle distribution of final particle stack. **(H)** Fourier shell correlation (FSC) versus resolution between half maps from final refinement for the 6-subunit map. **(I)** Final sharpened 7-subunit map colored by local resolution and viewed from the either side of the filament and from both the 3’ and 5’-ends. **(J)** Cutaway map showing resolution within the filament cavity of the 7-subunit map. **(K)** View angle distribution of final particle stack. **(L)** Fourier shell correlation (FSC) versus resolution between half maps from final refinement for the 7-subunit map.

**Fig. S23.**
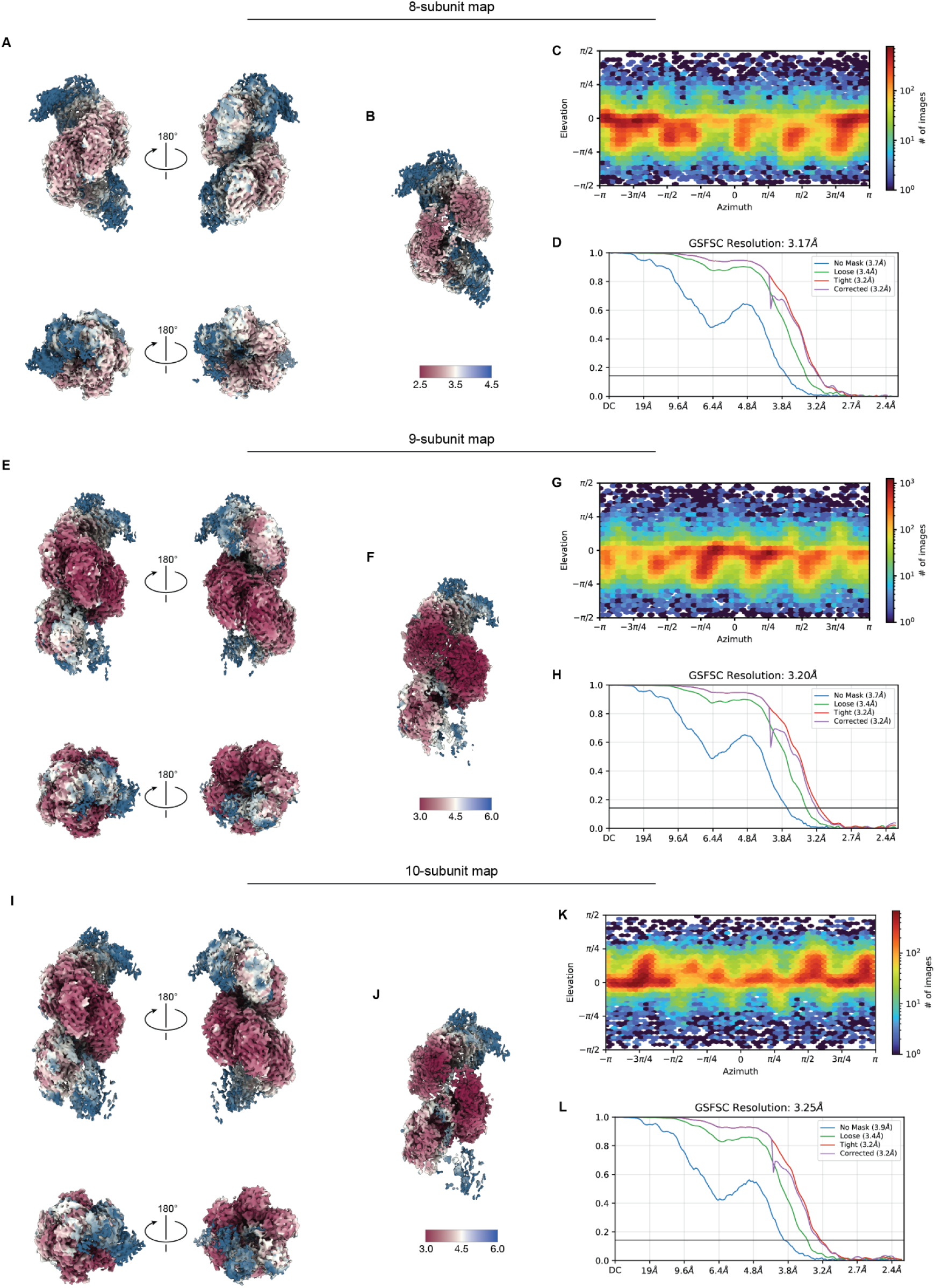
SUSP1 dsDNA ternary cryo-EM data validation for 8, 9, and 10 subunit conformations. **(A)** Final sharpened 8-subunit map colored by local resolution and viewed from the either side of the filament and from both the 3’ and 5’-ends. **(B)** Cutaway map showing resolution within the filament cavity of the 8-subunit map. **(C)** View angle distribution of final particle stack. **(D)** Fourier shell correlation (FSC) versus resolution between half maps from final refinement for the 8-subunit map. **(E)** Final sharpened 9-subunit map colored by local resolution and viewed from the either side of the filament and from both the 3’ and 5’-ends. **(F)** Cutaway map showing resolution within the filament cavity of the 9-subunit map. **(G)** View angle distribution of final particle stack. **(H)** Fourier shell correlation (FSC) versus resolution between half maps from final refinement for the 9-subunit map. **(I)** Final sharpened 10-subunit map colored by local resolution and viewed from the either side of the filament and from both the 3’ and 5’-ends. **(J)** Cutaway map showing resolution within the filament cavity of the 10-subunit map. (**K)** View angle distribution of final particle stack. **(L)** Fourier shell correlation (FSC) versus resolution between half maps from final refinement for the 10-subunit map.

**Fig. S24.**
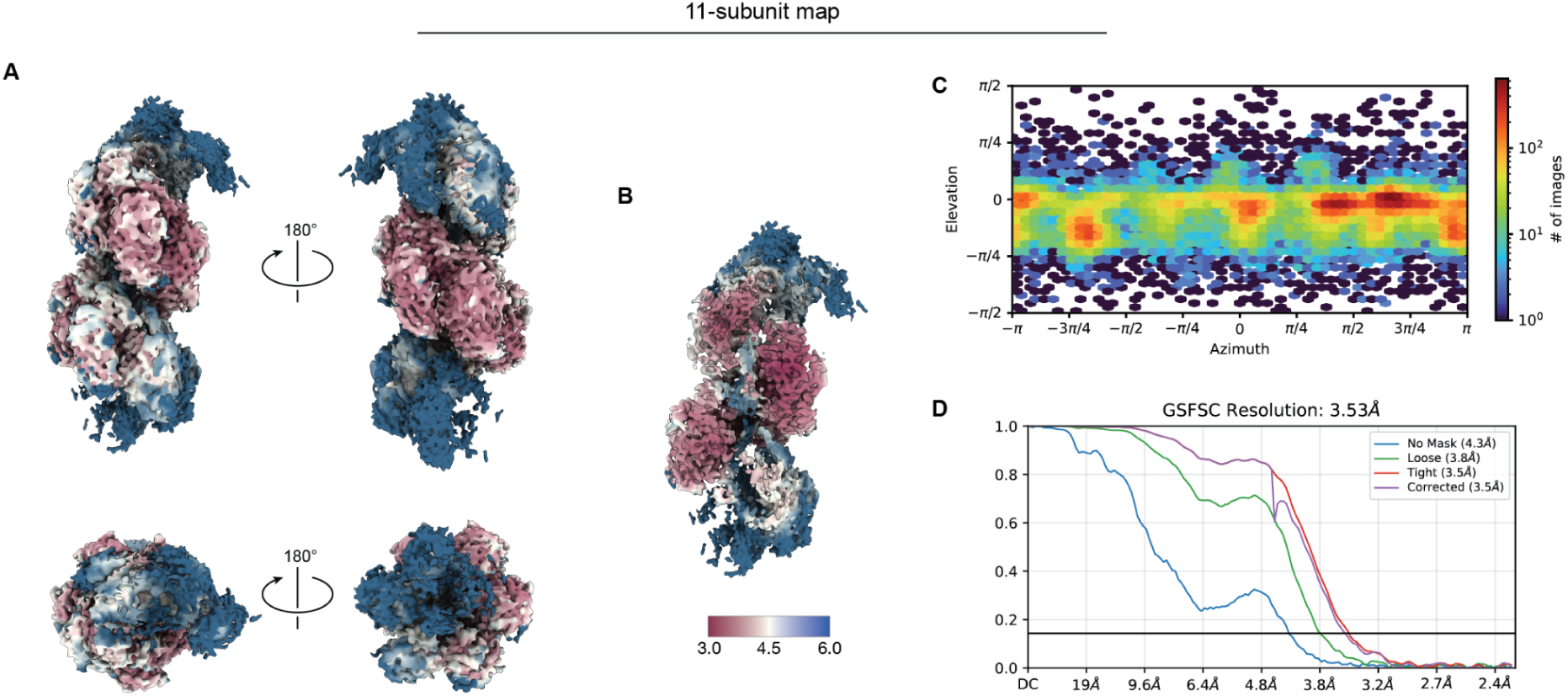
SUSP1 dsDNA ternary cryo-EM data validation for the 11 subunit conformation. **(A)** Final sharpened 11-subunit map colored by local resolution and viewed from the either side of the filament and from both the 3’ and 5’-ends. **(B)** Cutaway map showing resolution within the filament cavity of the 11-subunit map. **(C)** View angle distribution of final particle stack. **(D)** Fourier shell correlation (FSC) versus resolution between half maps from final refinement for the 11-subunit map.

**Fig. S25.**
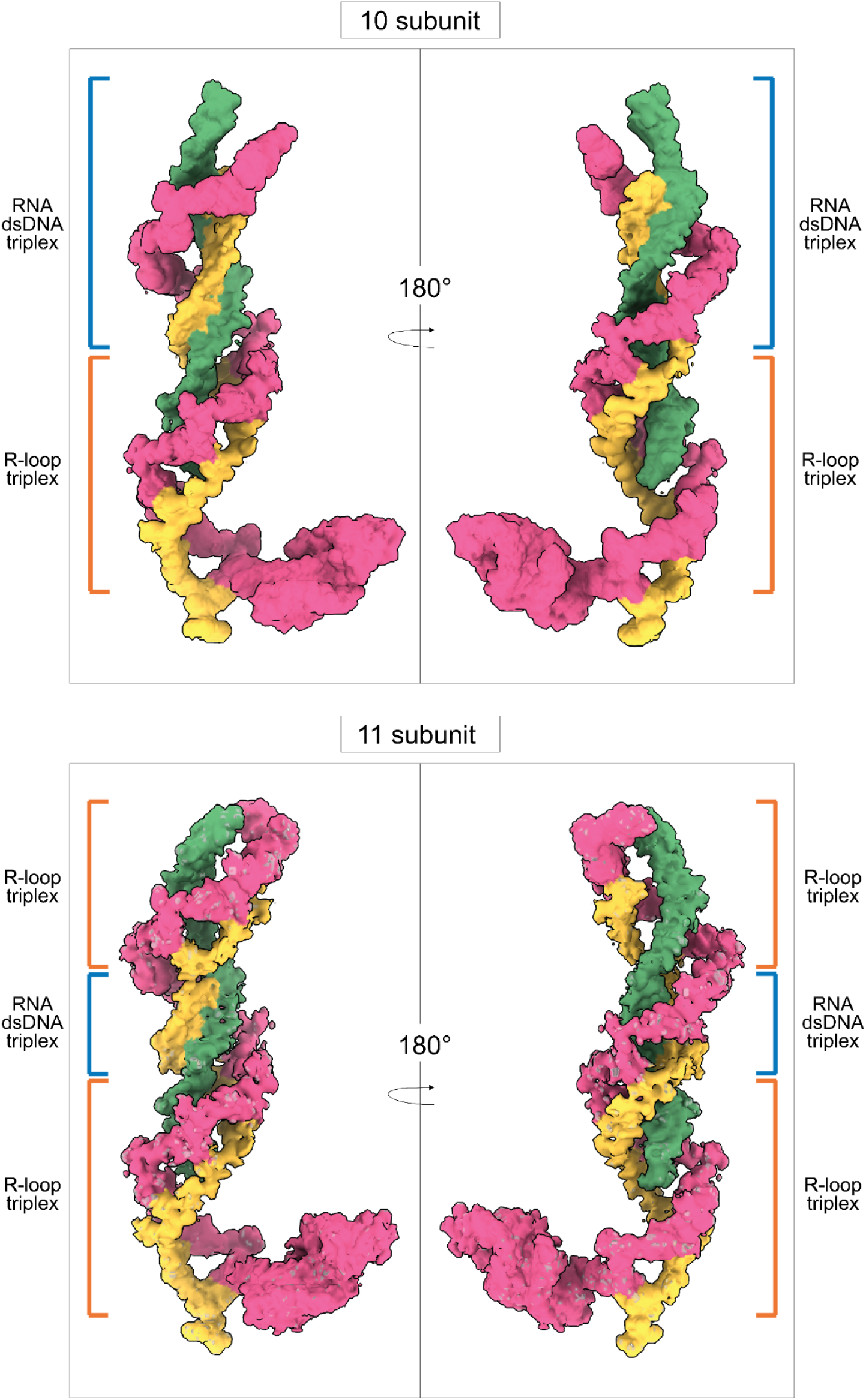
Comparison of SUSP1 dsDNA ternary complex with 10 and 11 resolved subunits. Carved nucleic acid densities of the 10– and 11-subunit states. The overall triplex architecture is preserved even when RNA-dsDNA to R-loop conversion occurs discontinuously along the filament.

**Table S1.** Sample preparation and freezing conditions.

| **Sample** | **VIPR**<br>**concentration** | **Target**<br>**Concentration** | **Sample Volume** | **Grid Type** |
| --- | --- | --- | --- | --- |
| SUSP1 Binary | 4 μM | - | 3.0 μl | UltrAuFoil 300<br>mesh R1.2/1.3 |
| SUSP1 VIPR +<br>pre-unwound<br>dsDNA | 1.3 μM | 2.6 μM | 3.0 μl | UltrAuFoil 300<br>mesh R1.2/1.3 |
| SUSP1 VIPR +<br>ssRNA | 2.25 μM | 5 μM | 3.0 μl | UltrAuFoil 300<br>mesh R1.2/1.3 |
| SUSP1 VIPR +<br>ssDNA | 1.3 μM | 2.6 μM | 3.0 μl | UltrAuFoil 300<br>mesh R1.2/1.3 |
| SUSP1 VIPR +<br>dsDNA | 1.3 μM | 2.6 μM | 3.0 μl | UltrAuFoil 300<br>mesh R1.2/1.3 |
| *P. fulva* VIPR | 4.5 μM | - | 3.5 μl | UltrAuFoil 300<br>mesh R1.2/1.3 |

**Table S2.**
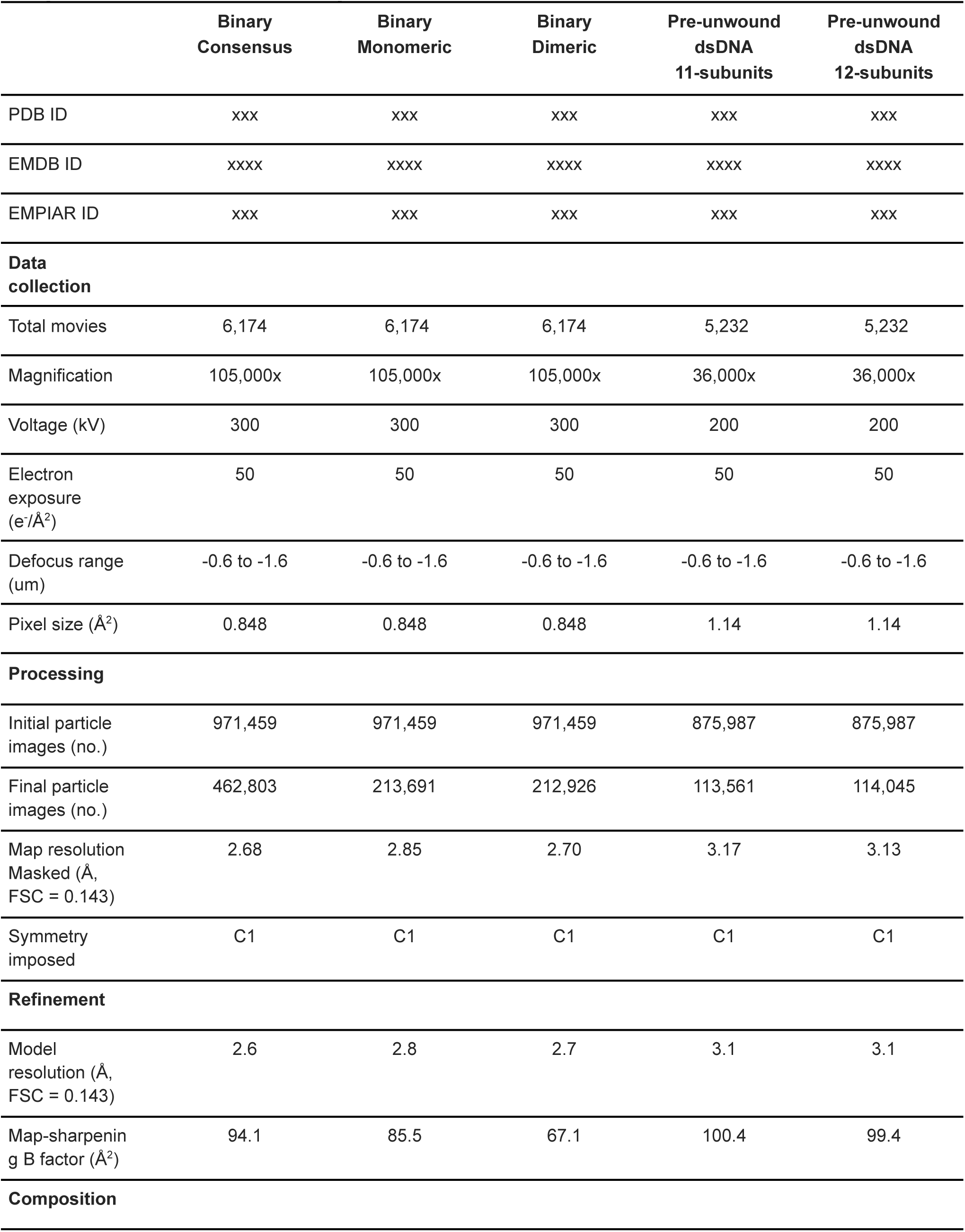
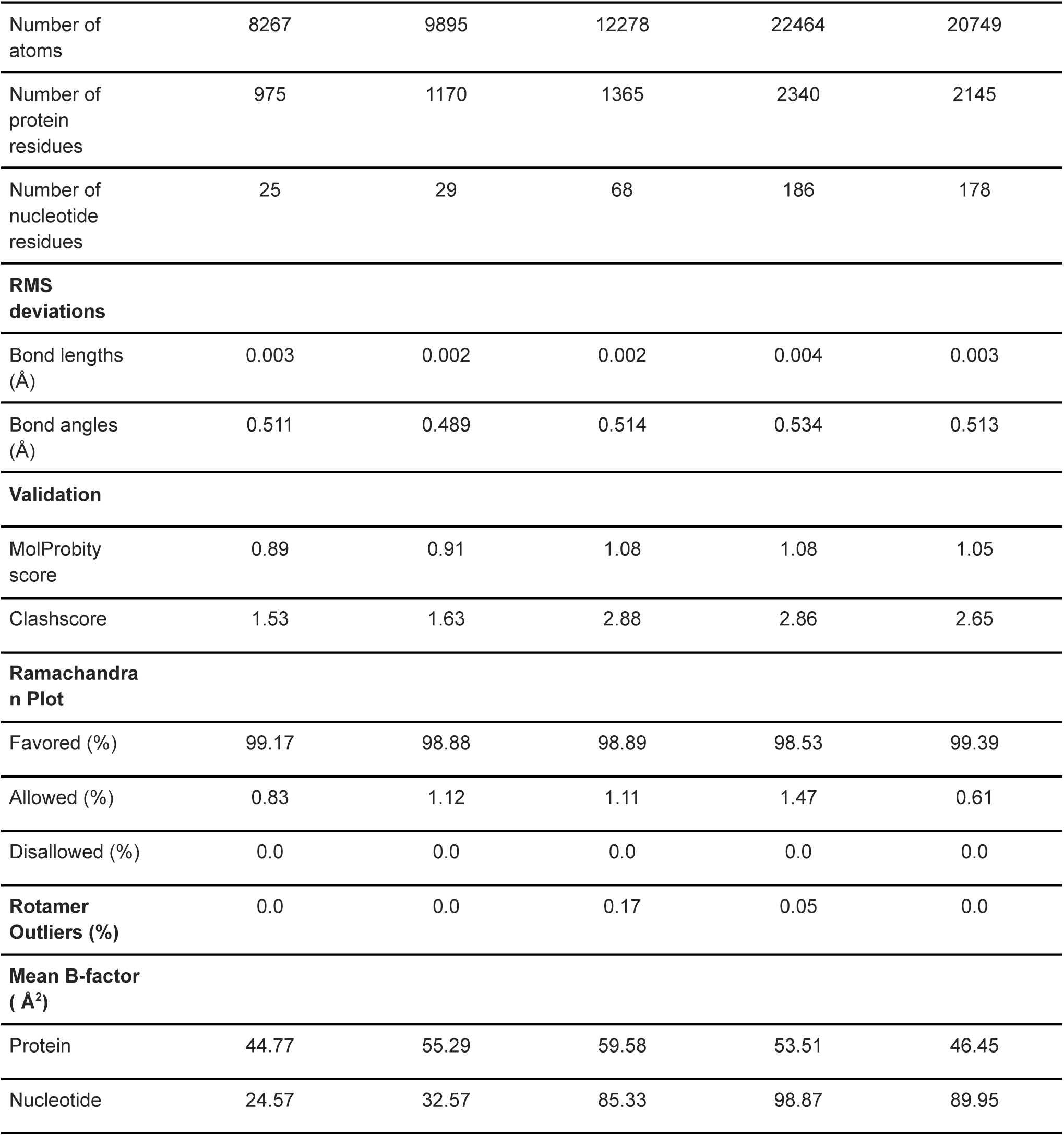
Cryo-EM data collection, refinement, and validation statistics for SUSP1 Binary and pre-unwound dsDNA Ternary.

**Table S3.**
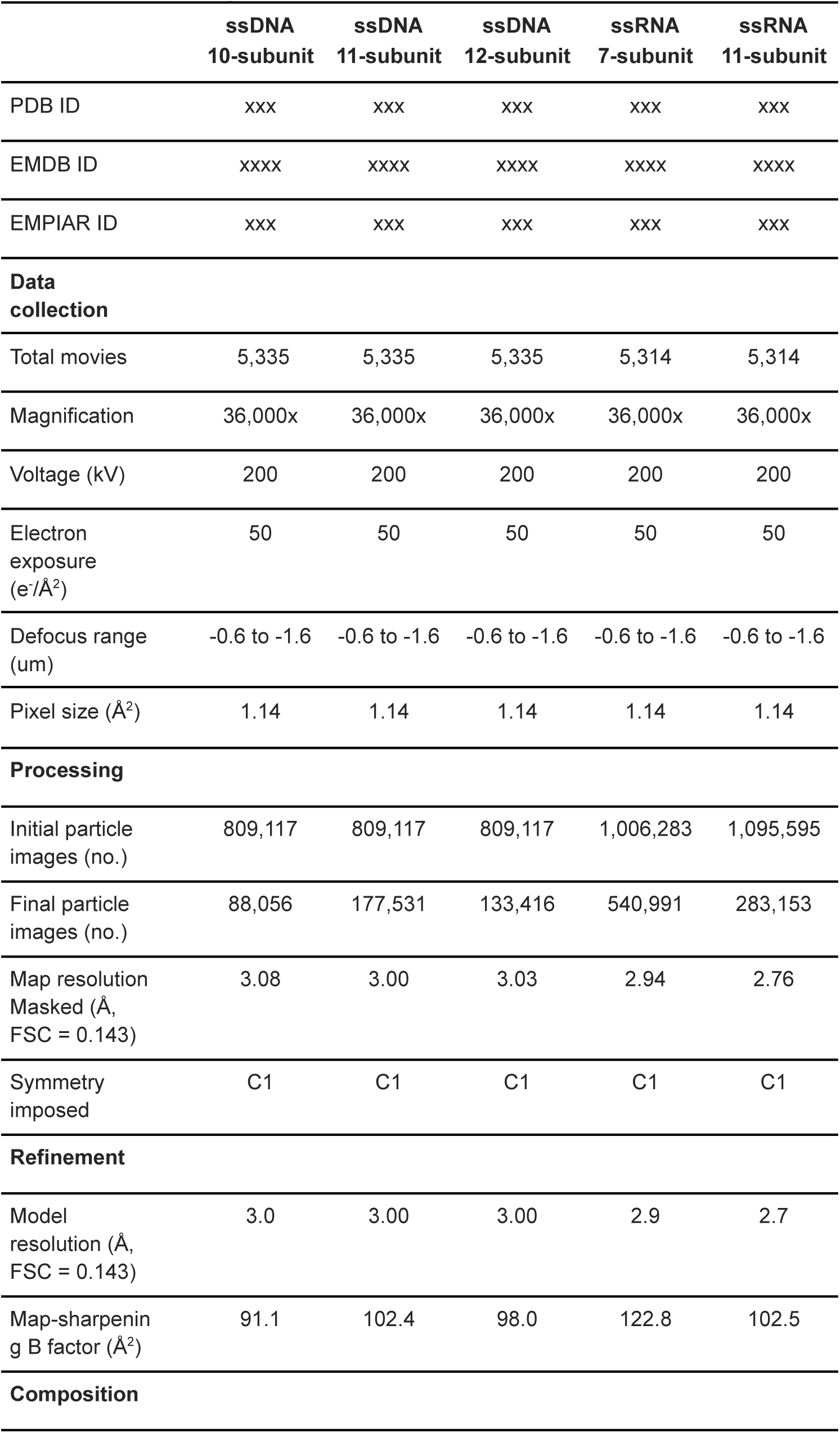
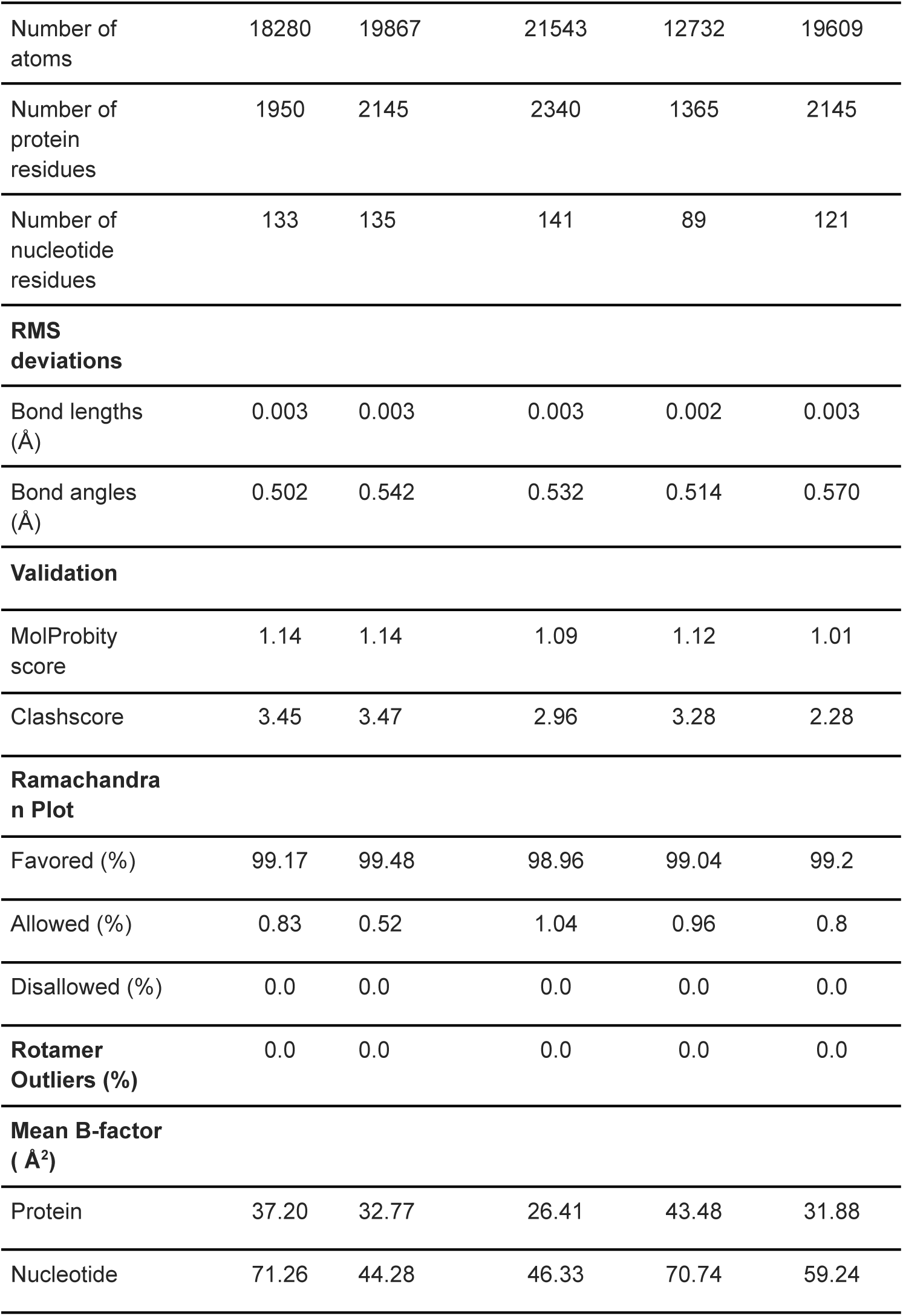
Cryo-EM data collection, refinement, and validation statistics for SUSP1 ssDNA and ssRNA Ternary.

**Table S4.**
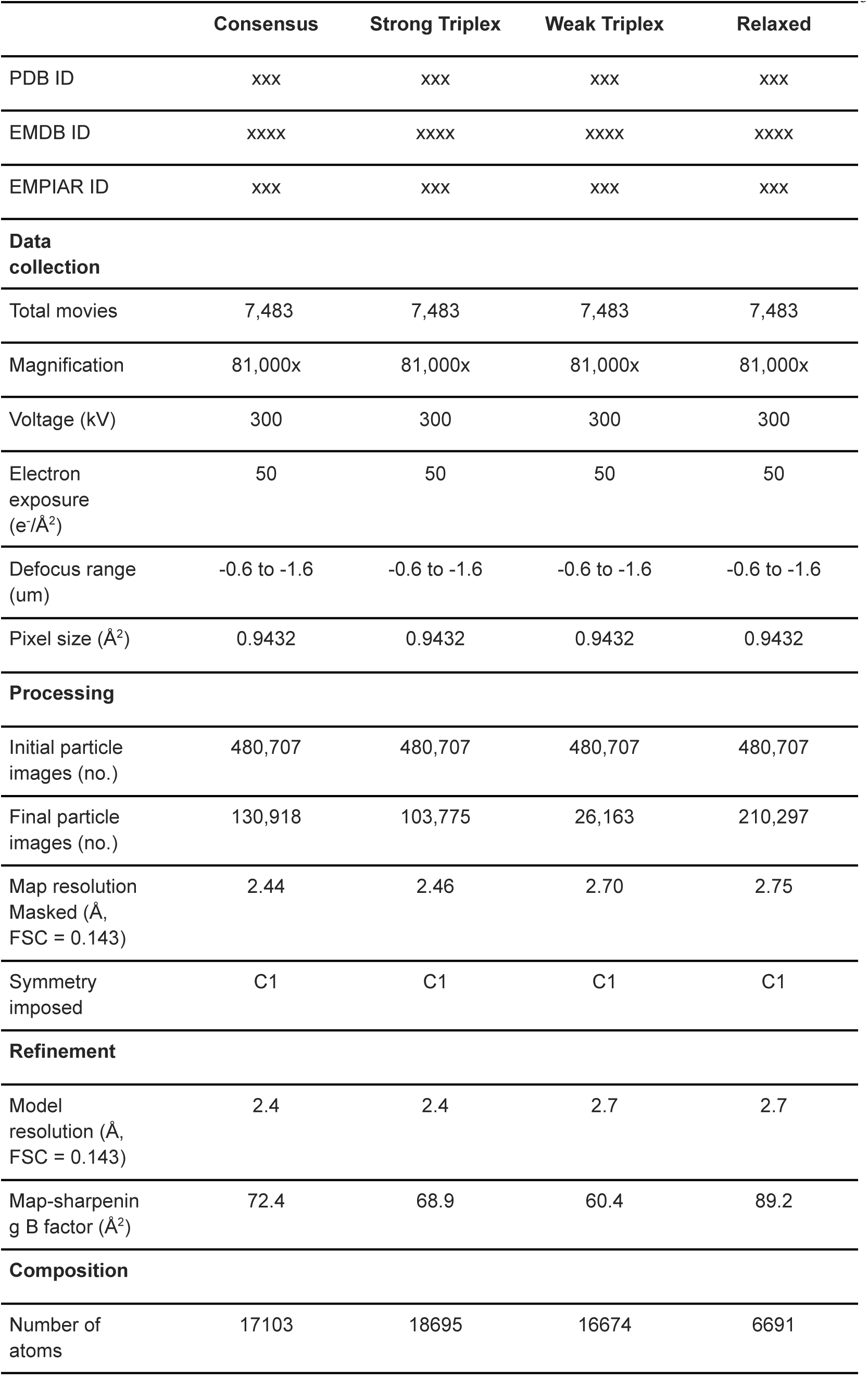
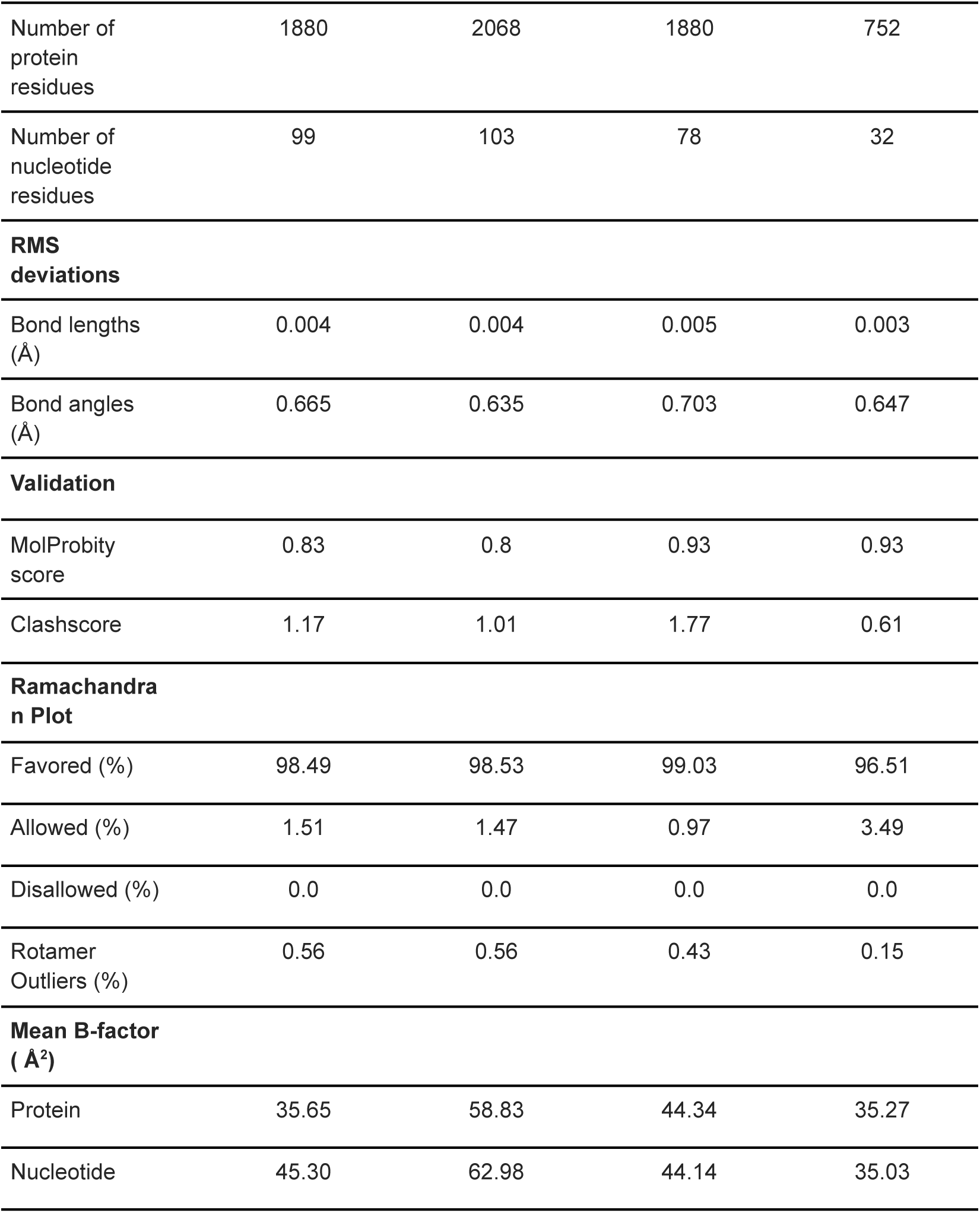
Cryo-EM data collection, refinement, and validation statistics for *P. fulva* VIPR.

**Table S5.** Cryo-EM data collection, refinement, and validation statistics for SUSP1 dsDNA Ternary.

|  | 5 subunit | 6 subunit | 7 subunit | 8 subunit | 9 subunit | 10 subunit | 11 subunit |
| --- | --- | --- | --- | --- | --- | --- | --- |
| PDB ID | xxx | xxx | xxx | xxx | xxx | xxx | xxx |
| EMDB ID | xxxx | xxxx | xxxx | xxxx | xxxx | xxxx | xxxx |
| EMPIAR ID | xxx | xxx | xxx | xxx | xxx | xxx | xxx |
| **Data collection** |  |  |  |  |  |  |  |
| Total movies | 9029 | 9029 | 9029 | 9029 | 9029 | 9029 | 9029 |
| Magnification | 36,000x | 36,000x | 36,000x | 36,000x | 36,000x | 36,000x | 36,000x |
| Voltage (kV) | 200 | 200 | 200 | 200 | 200 | 200 | 200 |
| Electron exposure (e<sup>-</sup>/Å<sup>2</sup>) | 50 | 50 | 50 | 50 | 50 | 50 | 50 |
| Defocus range (μm) | -0.6 to -1.6 | -0.6 to -1.6 | -0.6 to -1.6 | -0.6 to -1.6 | -0.6 to -1.6 | -0.6 to -1.6 | -0.6 to -1.6 |
| Pixel size (Å<sup>2</sup>) | 1.14 | 1.14 | 1.14 | 1.14 | 1.14 | 1.14 | 1.14 |
| **Processing** |  |  |  |  |  |  |  |
| Initial particle images (no.) | 1,271,801 | 1,271,801 | 1,271,801 | 1,271,801 | 1,271,801 | 1,271,801 | 1,271,801 |
| Final particle images (no.) | 33,174 | 134,829 | 185,176 | 133,189 | 155,696 | 115,442 | 49,155 |
| Map resolution Masked (Å, FSC = 0.143) | 3.35 | 3.25 | 3.17 | 3.17 | 3.20 | 3.25 | 3.53 |
| Symmetry imposed | C1 | C1 | C1 | C1 | C1 | C1 | C1 |
| **Refinement** |  |  |  |  |  |  |  |
| Model resolution (Å, FSC = 0.143) | 3.3 | 3.2 | 3.1 | 3.1 | 3.20 | 3.2 | 3.5 |
| Map-sharpening B factor (Å<sup>2</sup>) | 83.7 | 112.1 | 117.3 | 109.7 | 110.6 | 104.3 | 92.9 |
| **Composition** |  |  |  |  |  |  |  |
| Number of atoms | 9807 | 11331 | 13031 | 14702 | 16659 | 18503 | 20114 |
| Number of protein residues | 975 | 1170 | 1365 | 1560 | 1755 | 1950 | 2145 |
| Number of nucleotide residues | 98 | 97 | 104 | 110 | 130 | 144 | 147 |
| **RMS deviations** |  |  |  |  |  |  |  |
| Bond lengths (Å) | 0.003 | 0.003 | 0.003 | 0.004 | 0.003 | 0.003 | 0.002 |
| Bond angles (Å) | 0.508 | 0.648 | 0.560 | 0.554 | 0.578 | 0.583 | 0.527 |
| **Validation** |  |  |  |  |  |  |  |
| MolProbity score | 1.02 | 0.95 | 0.93 | 0.98 | 1.05 | 0.93 | 0.89 |
| Clashscore | 2.35 | 1.88 | 1.75 | 2.07 | 2.64 | 1.73 | 1.51 |
| **Ramachandra n Plot** |  |  |  |  |  |  |  |
| Favored (%) | 98.96 | 98.36 | 98.67 | 98.90 | 98.04 | 98.81 | 98.40 |
| Allowed (%) | 1.04 | 1.55 | 1.26 | 1.10 | 1.96 | 1.19 | 1.60 |
| Disallowed (%) | 0.0 | 0.0 | 0.7 | 0.0 | 0.0 | 0.0 | 0.0 |
| **Rotamer Outliers (%)** | 0.0 | 0.0 | 0.0 | 0.0 | 0.06 | 0.0 | 0.0 |
| **Mean B-factor ( Å²)** |  |  |  |  |  |  |  |
| Protein | 35.77 | 33.20 | 59.46 | 40.75 | 37.52 | 116.41 | 66.82 |
| Nucleotide | 86.81 | 88.66 | 92.85 | 135.01 | 74.83 | 149.79 | 134.02 |

